# From strength to stability: muscular holding capacity characterized by Adaptive Force and its potential relevance for musculoskeletal complaints and non-contact injuries

**DOI:** 10.1101/2025.09.08.670731

**Authors:** Laura V. Schaefer, Frank N. Bittmann

**Author notes:** Corresponding author: (LVS).

## Abstract

Musculoskeletal complaints and non-contact injuries remain poorly understood; conventional strength assessments show limited predictive value. They omit the function at which injuries occur—holding or braking under load. This is addressed by muscle stability quantified by Adaptive Force (AF)—the capacity to adapt to increasing loads while maintaining position. Available AF data were synthesized to characterize stable and unstable muscle states, evaluate parameter robustness, and explore practical relevance. 481 stable and 307 unstable trials of 112 muscles (elbow/hip flexors, pectoralis major) from six studies were pooled, supplemented by illustrative single cases. Both states were assessed in identical muscles by objectified manual muscle testing. AF parameters were compared by paired t-tests; the difference between states was regressed on five factors (muscle group, tester, sex, age, experiment) in linear mixed-effects models and two distribution-free tests. Robustness was examined with states regrouped by experimental condition, in leave-one-experiment-out, and blinded-tester analyses. Unstable muscles showed reduced maximal isometric AF (−84 N) but higher peak AF (+14 N). Their ratio (AF-Ratio) amounted to ∼56% in unstable and ∼99% in stable state (*d_z_*=2.7), with hardly any overlap; this was reproduced by condition-based grouping (agreement 97%, κ=0.94). A pre-defined AF-Ratio threshold (stable ≥98%) agreed similarly (97%, κ=0.93). Without pectoralis (n=3), no factor was associated with AF-Ratio difference; absolute parameters showed partial associations. In most cases, unstable muscles are not weak: they fail to access their full force capacity isometrically but generate it during lengthening. Muscles can switch instantly and reversibly between states in reaction to diverse stimuli, possibly based on shared neural substrates of motor control, somatosensation, and emotion. The functional instability syndrome (FIS) is proposed: muscle instability may destabilize joints and thus be a key factor in complaints and non-contact injuries. AF-Ratio is self-referenced and sensitive, suggesting applicability for screening, prevention, diagnostics, and AF-guided individualized therapy; its clinical value must be evaluated.

## Introduction

Musculoskeletal complaints and non-contact injuries remain among the most prevalent health issues in both athletic and general populations [1–6], yet their causal mechanisms are insufficiently understood. Non-contact injuries typically occur during active deceleration and dynamic stabilization—when external forces must be absorbed [7–16]. Paradoxically, muscle function is almost exclusively assessed during pushing actions (maximal voluntary isometric contraction (MVIC), isokinetic testing). However, converging evidence indicates that conventionally assessed muscle strength—whether absolute values, ratios or asymmetries—does not adequately explain the occurrence of musculoskeletal complaints and injuries: reviews have found that strength measures offer limited predictive validity for hamstring or ACL injuries [16–23], with none of 13 strength-related variables reaching significance in a meta-analysis of 78 prospective studies [18]; for leg, ankle and foot injuries, three of four prospective studies found hip muscle performance not to be a risk factor, and the review’s authors note that an observed association might result from the injury rather than contribute to it [24]. Existing systematic reviews may further overestimate the strength–injury association due to selection bias, as some restricted inclusion to studies reporting significant associations—yet even then, the reported evidence was only moderate [21]. Moreover, well-trained athletes without apparent strength deficits develop non-contact injuries and musculoskeletal complaints—which further questions the explanatory value of conventional strength parameters, considered alone, and points to the multifactorial etiology. In low back pain—one of the most prevalent musculoskeletal conditions—the picture is similar: lower-limb strength was reduced in patients for hip abductors, hip extensors and knee extensors but not for knee flexors, though the differences were small and hip strength was not related to the development of back pain [25]; moreover, there is no direct evidence that correcting strength imbalances by resistance or flexibility training reduces pain or improves function [26]. Collectively, these findings challenge the prevailing strength paradigm and raise the question whether the parameters typically examined capture the aspects of neuromuscular control that are functionally relevant for the development of complaints and injuries.

A critical, yet largely overlooked dimension of muscle function is the capacity to hold and adapt to external forces—rather than to produce force against a resistance. In everyday life and sport, muscles are frequently required to maintain or adjust joint positions against varying loads: during landing, changes of direction, or walking downstairs. These tasks demand a continuous adaptation of muscle tension to match the momentarily acting external force while preserving muscle length and joint position. This holding function fundamentally differs from pushing: it requires a permanent sensorimotor feedback loop in which proprioceptive and kinesthetic information about muscle length, muscle tension, joint angle, and external load is integrated, compared with the intended motor command, and continuously adjusted [27–29]. The limiting factor is therefore not maximal strength but the quality of sensorimotor control.

Recent research supports this distinction of holding (HIMA) and pushing isometric muscle actions (PIMA) which can be differentiated by objective parameters, with HIMA requiring more complex sensorimotor control [30–32]. However, most studies on HIMA have used constant loads. Adapting to variable, especially increasing external forces—as required in most functional activities—challenges the neuromuscular system even further.

The term muscle stability was introduced to describe this specific neuromuscular capacity: the ability of a muscle to maintain its length (isometric condition) while adapting to external forces. Unlike pushing strength, this capacity reflects the functional demands of everyday and sport activities more closely. If this capacity is compromised, joints may not be adequately stabilized, rendering the structures more vulnerable to strain—even under habitual loads. Muscle stability is thus the functional foundation upon which joint stability depends.

To quantify muscle stability, the concept of Adaptive Force (AF) was introduced [33,34] and further developed using both pneumatic measurement systems [35–38] and objectified manual muscle testing [39–45]. AF operationalizes this capacity against a continuously increasing external load and is thus based on HIMA. The maximal force achieved under isometric conditions (AFiso_max_) reflects the holding capacity. If AFiso_max_ approximates the peak force (AF_max_), the muscle is considered stable. If, however, the muscle begins to lengthen at submaximal intensities—yielding before reaching its force capacity—the muscle is classified as unstable; AFiso_max_ is then substantially lower than AF_max_. Crucially, this does not reflect a strength deficit: the peak force during the subsequent eccentric phase mostly equals or exceeds that under stable conditions or the MVIC. The muscle is not weak; it is unstable.

Previous studies have demonstrated that the holding capacity reacts with high sensitivity and immediacy to various stimuli. Disruptive inputs—such as negative emotional stimuli or proprioceptive irritation—led to an immediate and substantial reduction of AFiso_max_ (−44%) in healthy individuals [40–44]. Long COVID patients similarly showed markedly reduced muscle stability (−53%) in their disease state [45,46]. Conversely, supportive inputs—including positive emotional stimuli, proprioceptive re-adjustment, and targeted treatment—restored stability immediately [40–46]. Thus, muscles can switch instantaneously between stable and unstable states, strongly suggesting a reflexive mechanism. The neurophysiological pathways through which such diverse stimuli may affect motor control will be addressed in the Discussion.

The present study synthesizes and re-evaluates the available AF data from six previously published studies [40–45] by pooling 481 stable and 307 unstable trials from 112 muscles of 68 participants—with each muscle measured in both states within the same individual. The aims are to (1) characterize the fundamental differences between stable and unstable muscles based on AF parameters and (2) evaluate whether the key parameters are robust across potentially influencing factors. Single cases complement the analysis to illustrate, explore, and discuss practical relevance. The underlying hypothesis is that shifting the perspective from strength to muscle stability may provide a more adequate framework for understanding musculoskeletal complaints and non-contact injuries.

## Materials and methods

The present work is a pooled secondary analysis of all available AF data from six previously published studies with healthy participants [40–44] and long COVID patients [45], complemented by illustrative, hypothesis-generating single cases. AF was compared for identical muscles between their stable and unstable state. All AF measurements were performed at the Neuromechanics Laboratory of the University of Potsdam using a manual muscle test (MMT) objectified by a wireless handheld device (see 2.2). The MMTs were executed by the same two experienced testers in all six experiments (tester A: male, ∼64 yrs., 185 cm, 87 kg, MMT experience: ∼27 yrs.; tester B: female, ∼35 yrs., 168 cm, 55 kg, MMT experience: ∼8 yrs.). The measurement procedure and the data processing were identical across all experiments; the experiments differed only in the stimuli applied between measurements.

### Participants

A total of 112 muscles from 68 participants (31 m, 37 f; age: 33.9 ± 13.1 yrs., range: 19–70; body mass: 72.4 ± 12.3 kg, body height: 175.0 ± 9.9 cm; body mass and height available for 67/68 participants) were included in the present evaluation. Fifty-six of these muscles were elbow flexors, 53 hip flexors and three pectoralis major muscles (all from male participants). Forty-four participants contributed two muscles each; among them, three participants took part in two of the pooled experiments and are therefore counted as one participant each, resulting in 68 participants with 112 muscles in total. Fifty-one participants were healthy (i.e., asymptomatic; 28 m, 23 f; 30.1 ± 10.6 yrs., range: 19–70) and participated in the experiments designed to provoke instability or stability through emotional imagery [42,43], odor perception [44], or muscle spindle conditioning [40,41]. The remaining 17 participants had the medical diagnosis long COVID (3 m, 14 f; 45.2 ± 13.7 yrs., range: 21–63), in whom AF was measured in the disease state and after recovery [45]. Four additional participants (2 m, 2 f) served as illustrative single cases in specific contexts. They were not included in the inferential statistics to preserve the systematic character of the group analysis.

The studies on which this article is based were conducted according to the guidelines of the Declaration of Helsinki and were approved by the Ethics Committee of the University of Potsdam, Germany (protocol code 35/2018; amendment for long COVID: 70/2021). All participants gave their written informed consent to participate, including consent to the scientific use of their anonymized data. As the present work is a secondary analysis of already collected and anonymized data covered by this consent, no additional ethics approval was necessary.

### Technical equipment

The handheld device to objectify the MMTs records reaction force and limb kinematics simultaneously during the test, thereby providing continuous quantitative data (development funded by the German Federal Ministry for Economic Affairs and Energy; no. ZF4526901TS7). It contains strain gauges (Sourcing map, model: a14071900ux0076, precision: 1.0 ± 0.1%, sensitivity: 0.3 mV/V) and kinematic sensors (Bosch BNO055, 9-axis absolute orientation sensor, sensitivity: ± 1%) at a sampling rate of 180 Hz. Data were AD-converted and transmitted via Bluetooth to a tablet with measurement software (Sticky Notes, StatConsult, Magdeburg, Germany). The force profiles of this instrumented MMT have been shown to be reproducible, both within testers (ICC(3,1) = 0.992 and 0.995) and between the two testers (ICC(3,1) = 0.989) [39].

### Manual muscle tests and settings

AF was assessed during objectified standardized MMTs performed as break tests [40–45]: the tester applied a standardized increasing force (force profile as defined in [39]) on the participant’s limb by pushing against it in the direction of muscle lengthening, while the participant was instructed to maintain the starting position (HIMA). Hence, they had to adapt to the increasing force in a static position. All MMTs were performed in the supine position. Joint angles were standardized at 90° for elbow and hip flexor tests. For pectoralis, the shoulder was positioned in 90° forward flexion with full elbow extension and maximal internal rotation. The handheld device was positioned between the tester’s palm and the participant’s limb (distal forearm or thigh), with contact points marked for intra-individual standardization across trials.

The tester rated each trial as ‘stable’ (position maintained throughout the force increase—reflecting adequate adaptation), ‘unstable’ (limb yielding at submaximal intensities—inadequate adaptation; during the eccentric phase, the force typically continues to increase until peak force is reached) or ‘borderline’ if neither applied clearly. The rating was based solely on the tester’s perception during the test; all objective parameters were computed afterwards from the recorded signals, so testers had no access to them at the time of testing. In two experiments (odors [44], second imagery study [42]), testers were additionally blinded to the experimental condition. For details of settings and procedures see [40–45].

### Procedure

Although the AF assessment was identical across the six studies included (same testers, setting, device), the experiments designed to provoke muscle instability or stability differed: two studies compared unpleasant vs. pleasant food imagery [42,43], one investigated unpleasant vs. pleasant odors [44], two provoked a muscle spindle slack vs. physiological adjustment [40,41], and one measured AF in long COVID patients across disease state, after treatment, and recovery [45]. In all studies two to three trials were performed per muscle and condition. Three studies of healthy participants [40,42,44] included additional baseline measurements; in the first imagery study [43] ten baseline measurements were recorded but not reported in the publication.

Inclusion criterion for the present analysis was the availability of clear stable and unstable trials for at least one muscle per participant, regardless of the experimental condition. Classification into stable and unstable trials was based on the testers’ clinical ratings during the respective studies. The agreement of this classification with the a priori experimental hypotheses (unstable for negative imagery/odor, spindle slack, disease state; stable in the other condition) and with an objective classification based on a pre-specified AF-Ratio threshold is reported in the Results.

Of 801 trials originally included in the underlying studies, 23 were not considered in the present analysis: all five trials of one participant in whom only stable trials were available (Spindle Procedure), precluding a within-muscle comparison, all eight trials of one hip flexor muscle with a high proportion of borderline ratings (Long COVID), one trial because of incorrect execution (initial pushing of participant), and nine because the muscle state could not be clearly assigned (six ‘borderline’, three marked as ‘clearly more suspension’ during the test, although rated ‘stable’). With the ten additional baseline trials of the first imagery study, 788 trials from 112 muscles were available and included in the present evaluation. In 2.9% of the included trials (n = 23), the observed muscle state did not match the original hypothesis of the experimental condition (6 stable despite hypothesized instability, 17 vice versa); these were classified according to the observed state into stable or unstable (tester’s rating). This results in a total of 481 stable trials (238 elbow flexors, 232 hip flexors, 11 pectoralis) and 307 unstable trials (150 elbow flexors, 148 hip flexors, 9 pectoralis) for the statistical analysis.

### Data processing

The data were processed in the underlying studies; the corresponding values were used for the present analysis. Processing was performed using NI DIAdem 2017 (National Instruments, Austin, TX, USA) based on force and gyrometer signals, independently of the tester’s rating (for details see [40–45]). The following parameters were extracted (Fig 1):

- AF_max_ (N): peak force of the entire trial, reached either under isometric or eccentric conditions.
- AFiso_max_ (N): highest force value under isometric conditions, determined by the last zero crossing of the gyrometer signal before continuous limb movement (break point) indicating yielding. The corresponding force value refers to AFiso_max_. If the isometric position was maintained throughout: AFiso_max_ = AF_max_.
- AFosc (N): force at the onset of clear mechanical oscillations in force signal (≥ 4 consecutive maxima with dx < 0.15 s). Without such an oscillatory onset: AFosc = AF_max_. In two stable and one unstable trial, AFosc could not be determined; these trials were omitted from the corresponding muscle means, each of which still rested on at least one further trial, so that all 112 muscles contribute to every parameter.

**Fig 1.**
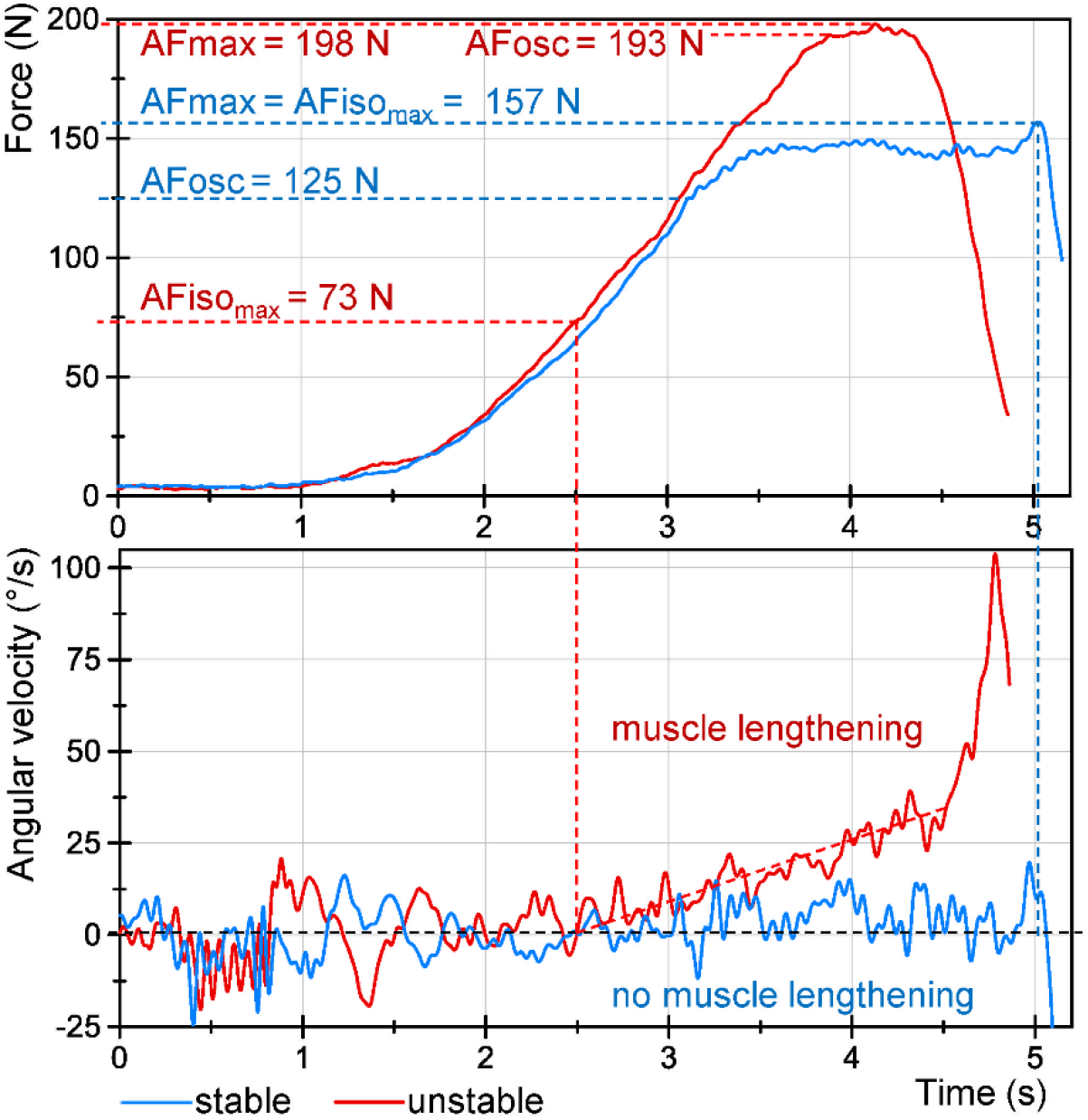
Representative stable vs. unstable AF trial of the same muscle. Displayed are the signals of force (upper panel) and angular velocity (gyro; lower panel) of two AF trials of elbow flexors of one healthy female participant (24 yrs.) during food imagery tested by the female tester. For pleasant imagery, the muscle maintained quasi-static position throughout (stable, blue), with AFiso_max_ = AF_max_. In contrast, for unpleasant imagery the muscle began to lengthen at a clearly submaximal force level (unstable, red), yet reached a high peak force during the eccentric phase, with AFiso_max_ substantially lower than AF_max_. In the stable trial, oscillations arose in the force signal at submaximal level; for the unstable trial, oscillations remained absent until near the peak force. The slopes of force increase were nearly identical for both stable and unstable trials, mirroring the standardized force application. Marked are the values of peak Adaptive Force (AF_max_), max. isometric AF (AFiso_max_) and AF at the onset of oscillations (AFosc) (all in N). Signals were filtered (Butterworth, low-pass, cut-off: 20 Hz, filter degree 5).

A conversion to torque was not performed. Within a muscle, the lever arm was identical across trials, so the comparisons between muscle states are unaffected. The ratios AFiso_max_/AF_max_ (AF-Ratio, given in %) and AFosc/AFiso_max_ were computed per trial and then averaged per muscle and state. AF-Ratio is self-referenced, relating the maximal holding capacity to the peak force of the same trial, and is therefore independent of absolute force level and lever arm; it serves as the main stability parameter [47]. AFosc/AFiso_max_ indicates whether oscillations arose before (< 1) or after (> 1) a potential break point. The slope of force rise was controlled in the underlying studies and did not differ between muscle states, a prerequisite for comparing the AF parameters.

### Statistical analysis

For evaluating differences between muscle state (stable vs. unstable) and potential influencing factors (muscle group, tester, participant sex, age, experiment), statistical analyses were performed in Python 3.14.2 (Python Software Foundation, Beaverton, OR, USA) with statsmodels 0.15, scipy 1.18.1, numpy 2.5.3, pandas 3.0.6, matplotlib 3.11.2; descriptive statistics and verification of the main results were additionally computed in Microsoft Excel (Version 2608; Microsoft Corp., Redmond, WA, USA). Descriptive data are given as arithmetic means ± standard deviations (M ± SD) with 95%-confidence intervals (CI). The variability of the trials within one muscle and state was described by the coefficient of variation (CV = SD/M), averaged over muscles. Differences between muscle states are given as unstable minus stable, so that negative values indicate lower values in the unstable state; averaged differences are reported as mean difference (MD). The significance level was α = 0.05. Holm adjustment was applied to the set of tests that jointly answer one question; the caption of each table states which tests that set comprises. Effect sizes are reported as *d_z_*, with values of 0.2, 0.5, 0.8, and ≥1.3 regarded as small, medium, large, and very large [48]. For the factor models, partial η² was computed from the Wald F statistic as η²p = F·df1/(F·df1 + df2).

#### Unit of analysis

The unit of analysis was the muscle. For each of the 112 muscles, all available trials of a given muscle state were averaged, yielding one stable and one unstable value per muscle and parameter. Because 44 of the 68 participants contributed two muscles—for three of them the two muscles were measured in different experiments—the dependence of muscles within participants was accounted for as described below.

#### Distributional assumptions and transformations

Normality of the within-muscle differences was assessed by the Shapiro-Wilk test together with skewness, kurtosis and Q-Q plots. Since the analyses operate on within-muscle differences, homogeneity of variance between muscle states is not required. Because AFosc/AFiso_max_ was markedly right-skewed and unbounded above, it was additionally analyzed after logarithmic transformation and is described by medians and interquartile ranges.

#### Comparison of muscle states

For each of the five parameters (AF_max_, AFiso_max_, AFosc, AF-Ratio and AFosc/AFiso_max_), the difference between muscle states was estimated in a linear mixed-effects model with a random intercept at participant level, since 44 of the 68 participants contributed two muscles; the model yields the mean difference with its 95% CI while accounting for this clustering. The standard errors were checked in an ordinary least squares model with the covariance clustered on participant. As supplementary comparisons, values of stable and unstable muscles were compared by paired t-tests, which are equivalent to a linear mixed-effects model with muscle state as fixed effect and a random intercept for the individual muscle. Shapiro-Wilk tests indicated normality of the within-muscle differences for AF_max_, AFosc and AF-Ratio (*p* = 0.139–0.774), but not for AFiso_max_ (*p* = 0.002) and AFosc/AFiso_max_ (*p* < 0.001); Wilcoxon signed-rank tests were therefore computed for all five parameters alongside the t-tests (Q-Q plots in S1 Fig). 95% CI for *d_z_* were obtained by a cluster bootstrap resampling participants (5000 replicates), as the 112 muscles are not independent. *P*-values of the muscle state comparisons were adjusted according to Holm across the five parameters, both for the model estimates and for the supplementary tests. Associations between parameters at muscle level were quantified by Pearson correlation coefficients with 95% CI from the same cluster bootstrap over participants.

#### Classification of muscle state

The assignment of trials to the stable and unstable state was based on the testers’ ratings during the experiments and formed the basis of all main analyses. To examine whether the results depend on this judgement, two further classifications were derived. The condition-based classification assigned each trial a state according to the hypothesis of the experimental condition (unstable: unpleasant imagery/odor, spindle slack, disease state pre-treatment; stable: baseline of healthy participants, pleasant or neutral imagery/odor, adjusted muscle spindle, post-treatment/recovery); it was determined a priori and is independent of both the testers’ perception and the instrumented measurement. The algorithm-based classification rated a trial stable if AF-Ratio reached ≥98%, the threshold defined for stable adaptation in the odor study [44]; the four experiments published after that study [40–42,45] thus provide data independent of its derivation. Agreement between the classifications was quantified as percentage agreement and Cohen’s kappa, with 95% CI from a cluster bootstrap over participants, and interpreted according to [49]. All main analyses were repeated with trials grouped by the condition-based classification. Since the algorithm-based classification is derived from AF-Ratio itself, it was used only for the agreement analysis.

#### Potential influencing factors

Whether the difference between muscle states depended on muscle group, tester, participant sex or experiment was tested by regressing it per muscle on each factor in a linear mixed-effects model with a random intercept at participant level, corresponding to the state × factor interaction of a repeated-measures design; age was tested in the same way as a continuous predictor. For the force parameters the difference was expressed as the relative change per muscle [(unstable − stable)/stable × 100], because absolute force values are not comparable across muscle groups and participants (lever arm, individual force level); the dimensionless ratios were compared as differences (unstable − stable). *P*-values were Holm-adjusted within each parameter (five factors), as the parameters represent distinct constructs. Since only three pectoralis major muscles were available, all tests were repeated without this muscle group. Shapiro–Wilk gave no evidence against normal residuals for AF_max_, AFiso_max_ and AF-Ratio (*p* = 0.14–0.72), but clear evidence against it for AFosc and AFosc/AFiso_max_ (*p* < 0.001). Homogeneity of the residual variance across factor levels was assessed by Brown–Forsythe tests on the same residuals. It was not violated for AFiso_max_ and AF-Ratio (all *p* ≥ 0.10), but was for AF_max_ and AFosc across tester, participant sex and experiment (*p* < 0.01) and for AFosc/AFiso_max_ across tester and participant sex (*p* = 0.038 and 0.043). All single-factor tests were therefore repeated by two procedures that assume neither normally distributed nor homoscedastic residuals (S4 Table): permutation tests under the null of exchangeable participants, permuting the complete label vector of a participant as a block among participants contributing the same number of muscles, which preserves the unbalanced cluster sizes (20,000 permutations); and wild cluster bootstrap tests of the same omnibus contrast, with the null imposed, Rademacher weights and clusters at participant level (9999 replicates), since cluster-robust Wald tests reject too often when a factor is identified from few clusters per level. Both were Holm-adjusted as above.

The data were pooled from six experiments that differed both in how the muscle states were induced and in their sample composition, so the factors are not independently distributed—each experiment has its own profile of testers, sexes and ages, and all three pectoralis major muscles come from one experiment. S1 Table gives their distribution across experiments, their pairwise association (bias-corrected Cramér’s V for the categorical factors [50], η for age) and the variance inflation in the joint model [51]. The factors are strongly entangled—Cramér’s V reaches 0.74 and twelve cells of the pairwise cross-tabulations are empty—but no standard error is inflated by more than a factor of 1.70 (participant sex). A model with all five factors was therefore fitted in addition: the single-factor model asks whether a factor is associated with the difference at all, the joint model whether that association persists once the others are accounted for. Joint-model *p*-values are unadjusted, since the five terms are estimated within one model. An effect appearing in only one model is reported as such and not interpreted as established.

#### Sensitivity analyses

Robustness of the pooled result was examined in four ways (S3 Table): by leave-one-experiment-out analyses, excluding each of the six experiments in turn; by excluding subgroups defined by content, namely the long COVID cohort (diseased persons), both spindle experiments (mechanical conditioning), the imagery and odor experiments (emotionally valent stimuli) and the pectoralis major muscles (only n = 3); by analyzing the muscles of each tester separately; and by restricting the analysis to the two experiments in which testers were blinded to the experimental condition (second imagery and odor [42,44]). In addition, the association of muscle group with the difference in AF-Ratio was re-examined with each of the three pectoralis major muscles omitted in turn.

The anonymized trial-level data and the complete analysis script are openly available from Zenodo [52] and are additionally provided as S1 Dataset and S1 File. The large language model Claude (Anthropic) assisted with statistical analyses and data processing scripts.

## Results

### AF parameters in stable vs. unstable muscles

All five parameters differed significantly between muscle states (Table 1, Fig 2); an ordinary least squares model with standard errors clustered on participant gave the same result.

**Fig 2.**
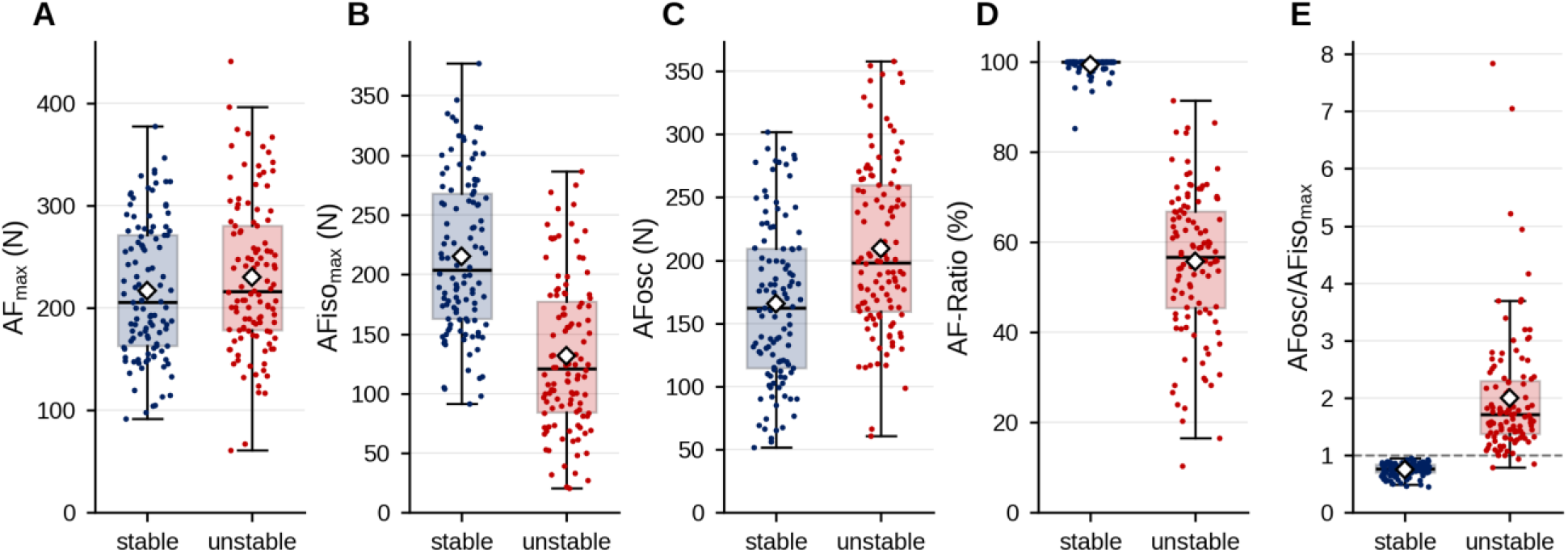
AF parameters of stable and unstable muscles. Values of all 112 muscles are shown as individual dots for the stable (blue) and unstable (red) state. The box contains the interquartile range (IQR, central 50% of muscles), the line inside the box marks the median, whiskers extend to the most extreme value within 1.5 × IQR, and white diamonds indicate arithmetic means. (**A**) peak Adaptive Force (AF_max_), (**B**) max. isometric AF (AFiso_max_), (**C**) AF at the onset of oscillations (AFosc), (**D**) AF-Ratio, (**E**) AFosc/AFiso_max_; the horizontal dashed line in (**E**) marks the value of 1, separating oscillations arising before (< 1) and after (> 1) the break point.

**Table 1.**
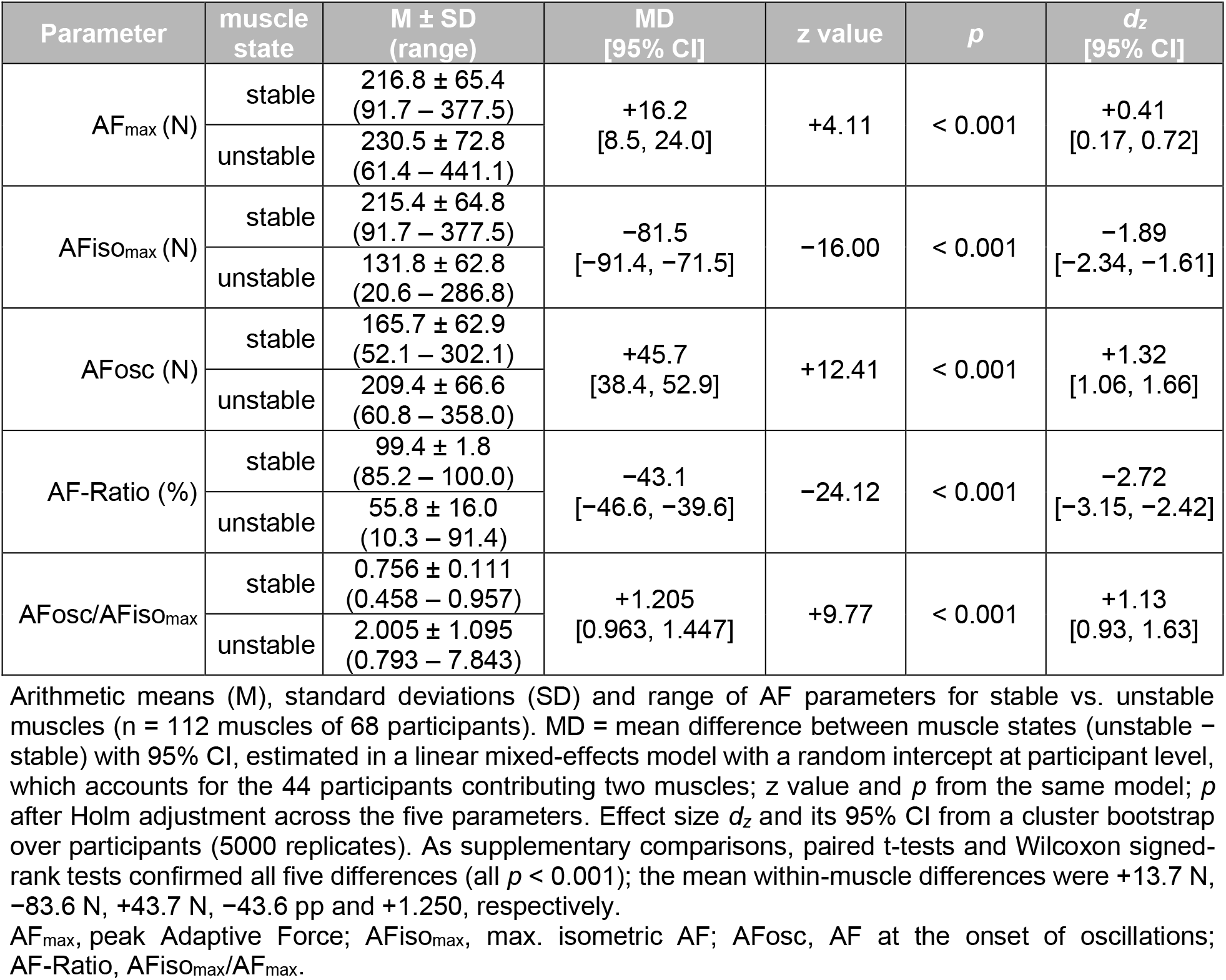
Parameters of Adaptive Force in stable vs. unstable muscles.

| **Parameter** | **muscle state** | **M ± SD<br>(range)** | **MD<br>[95% CI]** | **z value** | ***p*** | ***d*<sub>*z*</sub><br>[95% CI]** |
|---|---|---|---|---|---|---|
| AF<sub>max</sub> (N) | stable | 216.8 ± 65.4<br>(91.7 – 377.5) | +16.2<br>[8.5, 24.0] | +4.11 | < 0.001 | +0.41<br>[0.17, 0.72] |
|  | unstable | 230.5 ± 72.8<br>(61.4 – 441.1) |  |  |  |  |
| AFiso<sub>max</sub> (N) | stable | 215.4 ± 64.8<br>(91.7 – 377.5) | −81.5<br>[−91.4, −71.5] | −16.00 | < 0.001 | −1.89<br>[−2.34, −1.61] |
|  | unstable | 131.8 ± 62.8<br>(20.6 – 286.8) |  |  |  |  |
| AFosc (N) | stable | 165.7 ± 62.9<br>(52.1 – 302.1) | +45.7<br>[38.4, 52.9] | +12.41 | < 0.001 | +1.32<br>[1.06, 1.66] |
|  | unstable | 209.4 ± 66.6<br>(60.8 – 358.0) |  |  |  |  |
| AF-Ratio (%) | stable | 99.4 ± 1.8<br>(85.2 – 100.0) | −43.1<br>[−46.6, −39.6] | −24.12 | < 0.001 | −2.72<br>[−3.15, −2.42] |
|  | unstable | 55.8 ± 16.0<br>(10.3 – 91.4) |  |  |  |  |
| AFosc/AFiso<sub>max</sub> | stable | 0.756 ± 0.111<br>(0.458 – 0.957) | +1.205<br>[0.963, 1.447] | +9.77 | < 0.001 | +1.13<br>[0.93, 1.63] |
|  | unstable | 2.005 ± 1.095<br>(0.793 – 7.843) |  |  |  |  |

Arithmetic means (M), standard deviations (SD) and range of AF parameters for stable vs. unstable muscles (n = 112 muscles of 68 participants). MD = mean difference between muscle states (unstable − stable) with 95% CI, estimated in a linear mixed-effects model with a random intercept at participant level, which accounts for the 44 participants contributing two muscles; z value and *p* from the same model; *p* after Holm adjustment across the five parameters. Effect size *d<sub>z</sub>* and its 95% CI from a cluster bootstrap over participants (5000 replicates). As supplementary comparisons, paired t-tests and Wilcoxon signed-rank tests confirmed all five differences (all *p* < 0.001); the mean within-muscle differences were +13.7 N, −83.6 N, +43.7 N, −43.6 pp and +1.250, respectively.
AF<sub>max</sub>, peak Adaptive Force; AFiso<sub>max</sub>, max. isometric AF; AFosc, AF at the onset of oscillations; AF-Ratio, AFiso<sub>max</sub>/AF<sub>max</sub>.

AF_max_ was slightly higher in the unstable state (13.7 N, mean relative change per muscle: 7.7%, *d_z_* = 0.41), the only parameter without a large effect, reflected by the clear overlap of the distributions (Fig 2A). The ratio of unstable to stable AF_max_ amounted to 1.077 ± 0.202 (range 0.435–1.860), with 35 of 112 muscles showing a lower AF_max_ in the unstable state. AFiso_max_, in contrast, was 83.6 N lower in unstable muscles, a reduction of 39.8% (*d_z_* = −1.89); in 110 of 112 muscles AFiso_max_ was lower in the unstable state (ratio of unstable to stable AFiso_max_: 0.602 ± 0.197, range 0.092–1.141).

Both differences are combined in AF-Ratio, which showed the clearest distinction between muscle states: it was close to 100% in stable muscles (99.4 ± 1.8%) and dropped by 43.6 percentage points (pp) in the unstable state (*d_z_* = −2.72); the distributions of both states overlapped only between 85.2 and 91.4% (one stable and three unstable muscles; Fig 2D). AF-Ratio indicates that in the unstable state the static starting position could not be maintained up to the maximum force capacity during adaptation to the external force increase—AF_max_ was instead reached during the subsequent eccentric phase (muscle lengthening), at a level comparable to or even higher than in the stable state. In contrast, stable muscles reached their peak force under isometric conditions. Although AF_max_ forms the denominator of AF-Ratio, its higher value in the unstable state was not associated with the AF-Ratio difference (*r* = −0.06, *p* = 0.523); AF-Ratio thus varied with AFiso_max_, not with the peak force.

AFosc was reached at a 43.7 N higher force level in unstable muscles, 33.7% above the stable state (*d_z_* = 1.32). Accordingly, AFosc/AFiso_max_ was 1.250 higher in unstable muscles (*d_z_* = 1.13), below 1 in the stable and above 1 in the unstable state, i.e. oscillations arose before the break point in stable and after it in unstable muscles. On the log scale, which fits the skewed distribution better, the geometric means differed by a factor of 2.37 [2.18, 2.57] (*d_z_* = 1.96); given the skewness, the medians describe the distribution better than the means (stable: 0.769 [0.701, 0.845]; unstable: 1.709 [1.372, 2.301]). At trial level, oscillations arose before the break point in 469 of 479 stable trials (97.9%). Of the remaining ten, five had no detectable onset (AFosc = AF_max_) and five exceeded 1 (1.002–1.154). In unstable trials, oscillations arose after the break point in 280 of 306 trials (91.5%); 88 of these trials (28.8% of the 306 trials) showed no oscillatory onset at all (AFosc = AF_max_). In the remaining 26 unstable trials (8.5%), oscillations arose before AFiso_max_ (mean ratio 0.85).

The CV of the trials within one muscle and state was low for AF_max_ (stable 6.9 ± 5.1%, unstable 6.5 ± 4.7%) and for AF-Ratio in the stable state (1.0 ± 3.1%). In the unstable state, all parameters involving AFiso_max_ were markedly more variable (AFiso_max_ 22.7 ± 18.5%, AF-Ratio 21.9 ± 18.1%, AFosc/AFiso_max_ 25.3 ± 19.5%), whereas AFosc was not (stable 14.5 ± 7.3%, unstable 10.1 ± 9.4%).

### Distribution of the stability parameter AF-Ratio

At trial level, AF-Ratio of the two muscle states (tester rating) was distributed in different ranges, with few trials in between (Fig 3). In stable trials, it amounted to 99.5 ± 2.6% (median 100%, range 70.5–100%); 451 of 481 trials (94%) reached exactly 100%, i.e. AFiso_max_ was identical to AF_max_, and only 8 trials (1.7%) were below 90%, with no trial showing values below 70%. In unstable trials, it amounted to 56.4 ± 19.4% (median 58.1%, range 7.4–100%); only 1 of 307 trials (0.3%) reached 100%, while 299 trials (97%) were below 90%, 227 trials (74%) below 70%, and 112 trials (36%) even below 50%.

**Fig 3.**
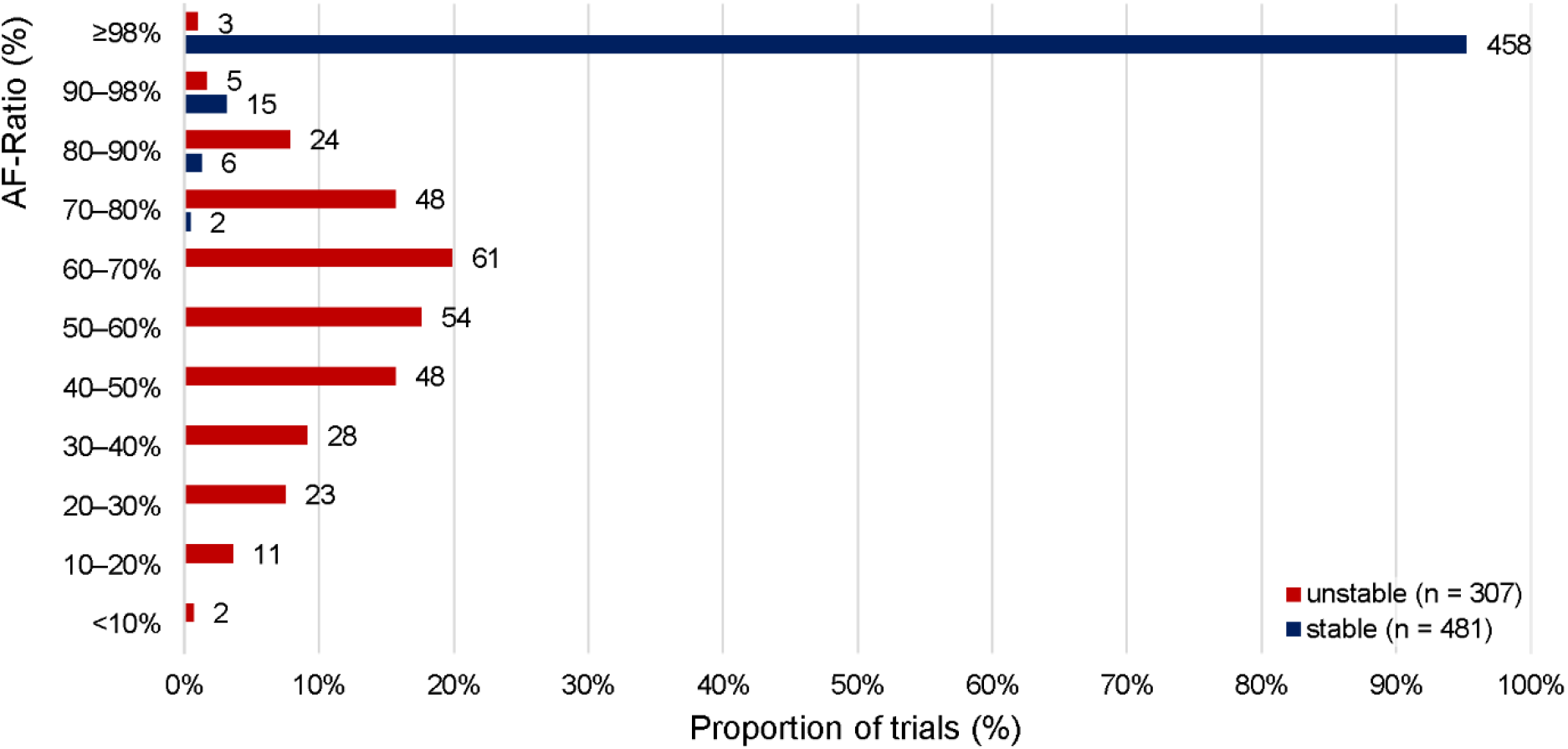
Distribution of AF-Ratio across stable and unstable single trials. Proportion of stable (blue, n = 481) and unstable trials (red, n = 307) within pre-defined ranges of AF-Ratio. The number of single trials is given beside each bar. The highest category corresponds to the threshold of 98% defined for stable adaptation in [44].

### Comparison of muscle state classification

All classifications were compared at trial level (n = 788). Applying the algorithm-based classification (stable = AF-Ratio ≥98%, pre-defined in [44]), 458 of 481 trials (95%) rated stable by the tester reached this threshold, and 304 of 307 trials (99%) rated unstable fell below it. Agreement between the testers’ ratings and this classification was 96.7% (κ = 0.93, 95% CI [0.90, 0.96]), corresponding to almost perfect agreement [49]. Restricted to the 643 trials of the four experiments published after the odor study [40–42,45], agreement was 97.4% (κ = 0.95 [0.92, 0.97]).

The condition-based classification agreed with the testers’ ratings in 765 of 788 trials (97.1%, κ = 0.94 [0.90, 0.97]). In 17 trials a muscle was rated unstable although a stable state had been hypothesized, in six trials the reverse occurred. Agreement between the condition-based and the algorithm-based classification, both independent of the testers’ perception, was 94.3% (κ = 0.88 [0.84, 0.92]).

At muscle level (n = 112), repeating the main comparison with trials grouped by the condition-based classification instead of the testers’ ratings yielded virtually unchanged results for all five parameters (AF_max_ MD = +11.9 N, *d_z_* = 0.38; AFiso_max_ MD = −84.1 N, *d_z_* = −1.87; AFosc MD = +42.7 N, *d_z_* = 1.32; AF-Ratio MD = −43.2 pp, *d_z_* = −2.51; AFosc/AFiso_max_ MD = +1.268, *d_z_* = 1.11; all *p* < 0.001).

### Potential influencing factors

Whether the difference between muscle states depended on muscle group, tester, participant sex, age or experiment was tested for each factor separately (Table 2, Fig 4, subgroup values in S2 Table). Since these factors were not independently distributed across participants—e.g., the Spindle Manipulation comprised exclusively young male football players, tester A measured mostly men, and the long COVID cohort was predominantly female, considerably older and the only cohort of diseased participants—an effect appearing in a single-factor model may belong to another factor. A model containing all five factors simultaneously was therefore fitted in addition; its results are reported at the end of this section. As distribution-free counterparts of the single-factor models, the permutation tests reproduced the decisions of Table 2 at the α = 0.05 level in 45 of the 50 comparisons and the wild cluster bootstrap in 44; all exceptions are marked there. For AF-Ratio, the permutation test agreed in all ten comparisons, and the bootstrap indicated no association with any of the five factors once the three pectoralis major muscles were excluded (*p* = 0.12–0.64).

**Fig 4.**
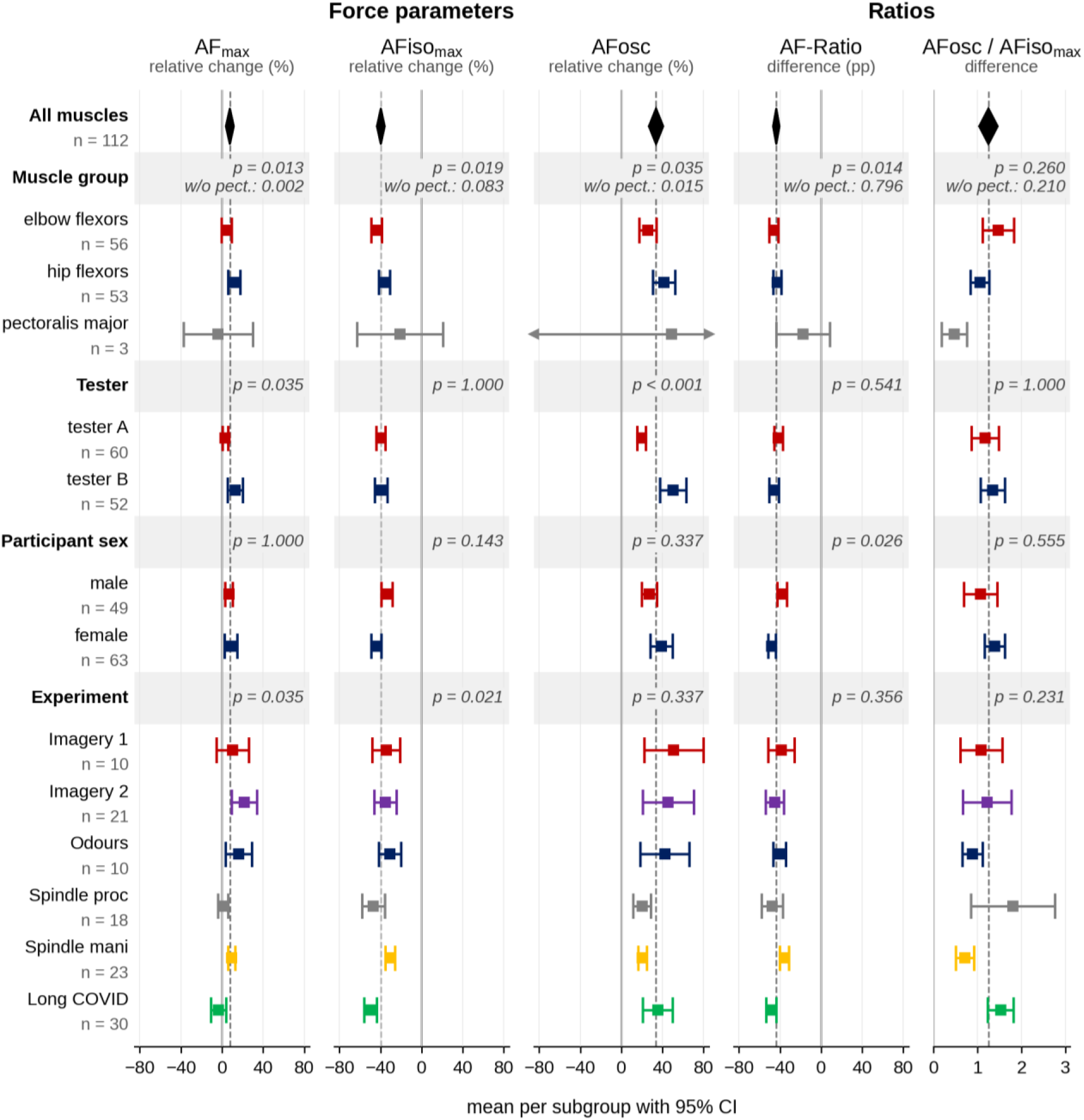
Forest plots of the difference between muscle states by potential influencing factor. Shown is the difference between the unstable and the stable state for the subgroups of the potential influencing factors muscle group, tester, participant sex and experiment; participant age, analyzed as a continuous covariate, is not displayed (all *p* ≥ 0.337). Force parameters are given as the mean relative change per muscle [(unstable − stable)/stable × 100], the ratios as the mean difference (unstable − stable; AF-Ratio in percentage points, pp). Squares show the subgroup mean with 95% CI, arrows indicate intervals extending beyond the displayed range; the diamonds and the dashed lines mark the mean across all 112 muscles. *P*-values are from the single-factor mixed models with a random intercept for participant, testing whether the differences depended on the subgroups of each factor, Holm-adjusted within each parameter across the five factors examined. Values for the pectoralis major are based on three muscles of three participants and are shown for completeness; they do not permit a meaningful group comparison. For muscle group, the *p*-value after exclusion of these muscles is therefore given in addition (w/o pect). AFiso_max_, maximal isometric Adaptive Force; AF_max_, peak AF; AFosc, AF at the onset of oscillations; AF-Ratio, AFiso_max_/AF_max_ × 100; *p*, *p*-value.

**Table 2.** Association of the potential influencing factors with the difference of AF parameters between the unstable and stable muscle states.

| **Model** | **Muscles** | **AF<sub>max</sub>** | **AFiso<sub>max</sub>** | **AFosc** | **AF-Ratio** | **AFosc/AFiso<sub>max</sub>** |
|---|---|---|---|---|---|---|
| **Muscle group** |  |  |  |  |  |  |
| single-factor | all | **0.013 (0.103)** | **0.019 (0.097)** | **0.035 (0.083)**<sup>a</sup> | **0.014 (0.103)**<sup>b</sup> | 0.260 (0.049) |
|  | w/o pect | **0.002 (0.112)** | 0.083 (0.049) | **0.015 (0.075)**<sup>a</sup> | 0.796 (0.012) | 0.210 (0.036) |
| joint | all | **0.006 (0.096)** | **0.009 (0.088)** | **0.017 (0.077)** | **0.007 (0.094)** | 0.144 (0.038) |
|  | w/o pect | **0.001 (0.102)** | 0.062 (0.035) | **0.015 (0.059)** | 0.458 (0.006) | 0.119 (0.024) |
| **Tester** |  |  |  |  |  |  |
| single-factor | all | **0.035 (0.059)**<sup>b</sup> | 1.000 (0.002) | **< 0.001 (0.149)** | 0.541 (0.011) | 1.000 (0.004) |
|  | w/o pect | 0.059 (0.050) | 1.000 (0.002) | **< 0.001 (0.133)** | 0.796 (0.010) | 1.000 (0.003) |
| joint | all | 0.081 (0.030) | 0.380 (0.008) | **< 0.001 (0.106)** | 0.817 (0.001) | 0.463 (0.005) |
|  | w/o pect | 0.264 (0.013) | 0.652 (0.002) | **0.010 (0.064)** | 0.743 (0.001) | 0.607 (0.003) |
| **Participant sex** |  |  |  |  |  |  |
| single-factor | all | 1.000 (0.003) | 0.143 (0.035) | 0.337 (0.022) | **0.026 (0.065)** | 0.555 (0.016) |
|  | w/o pect | 0.700 (0.002) | 0.282 (0.026) | 0.291 (0.025) | 0.119 (0.047) | 0.804 (0.011) |
| joint | all | 0.984 (0.000) | 0.289 (0.011) | 0.930 (0.000) | 0.266 (0.012) | 0.872 (0.000) |
|  | w/o pect | 0.713 (0.001) | 0.217 (0.015) | 0.787 (0.001) | 0.253 (0.013) | 0.954 (0.000) |
| **Age** |  |  |  |  |  |  |
| single-factor | all | 1.000 (0.004) | 1.000 (0.000) | 0.337 (0.023) | 0.738 (0.001) | 1.000 (0.000) |
|  | w/o pect | 0.700 (0.008) | 1.000 (0.002) | 0.291 (0.011) | 0.796 (0.001) | 1.000 (0.000) |
| joint | all | 0.310 (0.010) | **0.035 (0.043)** | 0.189 (0.017) | **0.039 (0.042)** | 0.133 (0.022) |
|  | w/o pect | 0.618 (0.003) | 0.102 (0.027) | 0.580 (0.003) | 0.062 (0.035) | 0.111 (0.026) |
| **Experiment** |  |  |  |  |  |  |
| single-factor | all | **0.035 (0.133)**<sup>a,b</sup> | **0.021 (0.144)**<sup>b</sup> | 0.337 (0.080) | 0.356 (0.078) | 0.231 (0.100) |
|  | w/o pect | **0.018 (0.150)**<sup>a,b</sup> | **0.026 (0.147)**<sup>a</sup> | 0.291 (0.085) | 0.300 (0.091) | 0.210 (0.104)<sup>b</sup> |
| joint | all | 0.066 (0.096) | **0.005 (0.149)** | 0.864 (0.018) | 0.071 (0.094) | **0.044 (0.105)** |
|  | w/o pect | 0.088 (0.091) | **0.013 (0.134)** | 0.961 (0.010) | 0.095 (0.089) | **0.041 (0.109)** |

For each parameter the difference between the two muscle states was modeled with a linear mixed model with a random intercept for participant—the relative change for AF<sub>max</sub>, AFiso<sub>max</sub> and AFosc, the difference for AF-Ratio and AFosc/AFiso<sub>max</sub>. Every factor was first tested in a model of its own (single-factor model) and then with all five factors entered simultaneously (joint model). Given is the *p*-value of the omnibus Wald test with the partial η² in parentheses; each model is reported for all 112 muscles (all) and after exclusion of the three pectoralis major muscles (w/o pect).
In the single-factor models *p* is Holm-adjusted within each parameter across the five factors. In the joint model the five terms are estimated within one model, and *p* is reported unadjusted. The partial η² of the two models are not directly comparable: in a single-factor model there is nothing to adjust for, whereas in the joint model η² refers to the variance remaining once the other four factors have been accounted for. Values with *p* < 0.05 are set in bold. The subgroup values underlying these tests are given in S2 Table.
<sup>a</sup> Not confirmed by the permutation test at the α = 0.05 level (S4 Table). The corresponding permutation *p*-values are: muscle group on AFosc 0.136 (all) and 0.056 (w/o pect); experiment on AF<sub>max</sub> 0.057 (all) and 0.063 (w/o pect); experiment on AFiso<sub>max</sub> 0.051 (w/o pect).
<sup>b</sup> Not confirmed by the wild cluster bootstrap at the α = 0.05 level (S4 Table). The corresponding bootstrap *p*-values are: tester on AF<sub>max</sub> 0.195 (all); experiment on AF<sub>max</sub> 0.076 (all) and 0.063 (w/o pect); experiment on AFiso<sub>max</sub> 0.062 (all); muscle group on AF-Ratio 0.356 (all); experiment on AFosc/AFiso<sub>max</sub> 0.029 (w/o pect)—the only case in which the bootstrap indicated an effect that the mixed model did not.
AFiso<sub>max</sub>, maximal isometric Adaptive Force; AF<sub>max</sub>, peak Adaptive Force; AFosc, Adaptive Force at the onset of oscillations; AF-Ratio, AFiso<sub>max</sub>/AF<sub>max</sub> × 100; *p*, *p*-value; η², partial eta squared; pect, pectoralis major muscles; w/o, without.

### Single-factor models

#### Muscle group

The difference between the unstable and stable muscle states depended on muscle group for all parameters except AFosc/AFiso_max_ (Table 2, Fig 4, S2 Table). For AF-Ratio this was attributable entirely to the three pectoralis major muscles: excluding them, elbow and hip flexors showed a similar drop in the unstable state (−46.1 vs. −42.6 pp, *p* = 0.796); the wild cluster bootstrap indicated no muscle-group effect on AF-Ratio even with these three muscles included (*p* = 0.356). The same applied to AFiso_max_, where the reduction tended to be larger in elbow than in hip flexors (−44.0% vs. −36.3%, *p* = 0.083). The differences between muscle groups for AF_max_ and AFosc persisted without the pectoralis major, with a higher relative change in hip than in elbow flexors (AF_max_: +11.8% vs. +4.4%, *p* = 0.002; AFosc: +41.5% vs. +25.6%, *p* = 0.015). For AFosc, the bootstrap confirmed the muscle-group effect (*p* = 0.042 and 0.011), whereas the permutation test did not (*p* = 0.136 for all muscles, *p* = 0.056 without pectoralis).

Fig 5 illustrates the distribution of AF-Ratio separated by muscle group and muscle state, reflecting the similar pattern especially for elbow and hip flexors. Notably, in the unstable state AF-Ratio dropped below 50% in 22 of 56 elbow flexors (39%) and 13 of 53 hip flexors (25%). The three pectoralis major muscles showed comparatively high values in the unstable state (82.2 ± 10.5%, range 70.8–91.4%). Variability across muscles was also clearly higher in the unstable vs. stable state in every muscle group (CV elbow 31.2% vs. 1.4%, hip 25.0% vs. 2.2%, pectoralis 12.7% vs. 0.0%).

**Fig 5.**
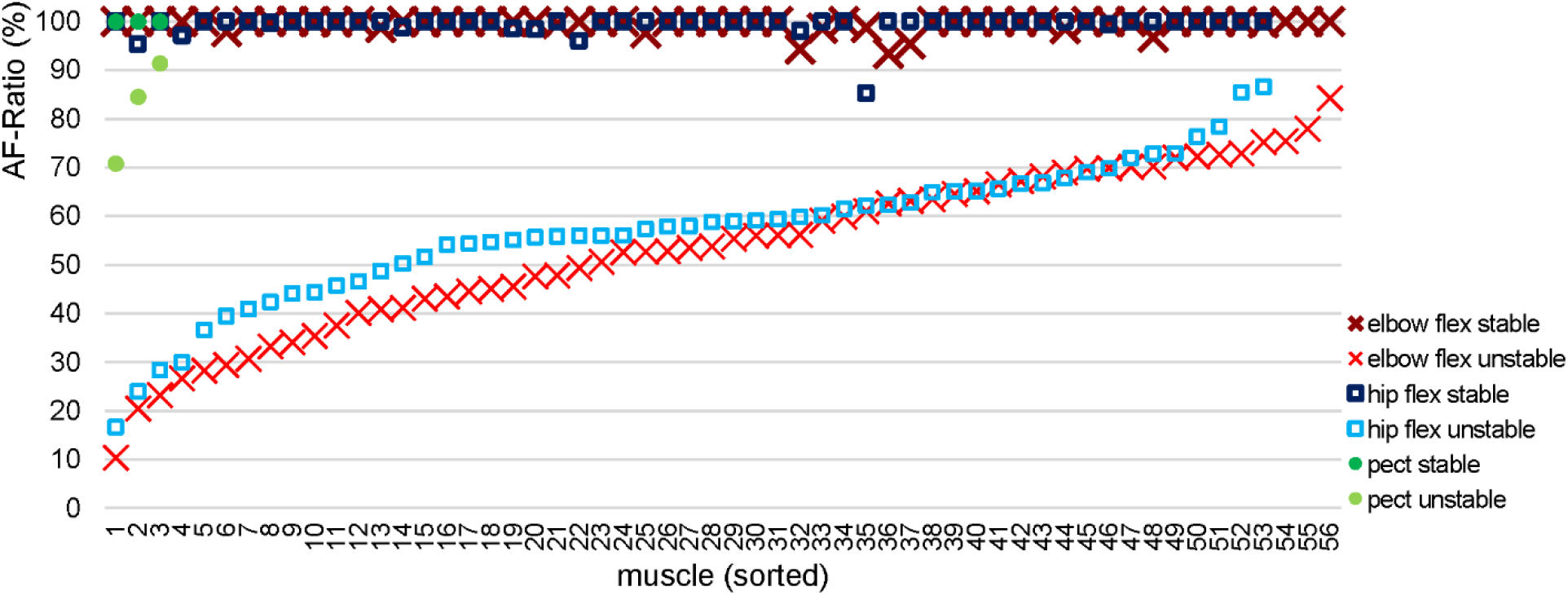
AF-Ratio per muscle, sorted in ascending order. Arithmetic mean of AF-Ratio per muscle (elbow flexors: red cross; hip flexors: blue square; pectoralis major: green circle; stable: dark, unstable: light). Each position on the x-axis represents one muscle, shown in both states; within each muscle group the muscles are sorted by their AF-Ratio in the unstable state. flex, flexors; pect, pectoralis major.

#### Tester

The relative change of AFiso_max_ between muscle states was practically identical for both testers (−39.9% vs. −39.6%, *p* = 1.000); AF-Ratio (−41.6 vs. −46.0 pp, *p* = 0.541) and AFosc/AFiso_max_ (1.169 vs. 1.343, *p* = 1.000) did not differ significantly between testers either (Table 2, Fig 4, S2 Table). Tester B showed a significantly higher relative change than tester A for AF_max_ (12.9% vs. 3.2%, *p* = 0.035) and AFosc (50.1% vs. 19.5%, *p* < 0.001); for AF_max_ this difference did not persist without the pectoralis major (*p* = 0.059). The bootstrap did not confirm the tester effect on AF_max_ (*p* = 0.195 for all muscles, *p* = 0.229 without pectoralis), whereas the effect on AFosc was confirmed throughout (*p* ≤ 0.002).

#### Participant sex

Participant sex affected AF-Ratio only (*p* = 0.026): the drop between the unstable and stable muscle states was significantly larger in females than in males (−48.0 vs. −38.0 pp). The difference arose in the unstable state, where females reached even lower values than males (51.6 ± 14.8% vs. 61.2 ± 16.0%), while the values in the stable state were nearly identical (99.6 ± 1.0% vs. 99.2 ± 2.5%); however, this effect did not persist when the pectoralis major was excluded (*p* = 0.119). For the other four parameters, no effect of sex was found (all *p* ≥ 0.143; Table 2, Fig 4, S2 Table).

#### Age

Participant age showed no association with the difference between muscle states in any parameter (all *p* ≥ 0.337, Table 2); the confidence intervals of all slopes included zero (S2 Table).

#### Experiment

For AF_max_ and AFiso_max_, the relative change between muscle states depended on the experiment (*p* = 0.035 and *p* = 0.021); for AFosc, AF-Ratio and AFosc/AFiso_max_ no effect was found (all *p* ≥ 0.231). Excluding the three pectoralis major muscles did not alter this (Table 2, Fig 4). In the distribution-free checks both effects were largely marginal: for AF_max_ neither the permutation test (*p* = 0.057 and 0.063) nor the bootstrap (*p* = 0.076 and 0.063) confirmed it in either sample, and for AFiso_max_ each confirmed it in one of the two samples (permutation *p* = 0.043 for all muscles, bootstrap *p* = 0.047 without pectoralis). The bootstrap additionally indicated an effect on AFosc/AFiso_max_ without the pectoralis major (*p* = 0.029).

For AFiso_max_ the experiments differed in extent, not in direction: all six showed clearly lower values in the unstable compared to the stable state, from −50.0% (Long COVID) and −47.2% (Spindle Procedure) to −30.5% (Spindle Manipulation), all significant in the MD-based within-experiment comparison (*p* ≤ 0.001; S2 Table).

For AF_max_ the picture was less uniform, and the two experiments with the largest AFiso_max_ reduction in the unstable state were those in which AF_max_ did not change considerably: Long COVID and Spindle Procedure (−3.2% and +1.1%, both *p* = 1.000 in the within-experiment comparison); the other four experiments showed a higher AF_max_ in the unstable state, ranging from +9.5% (Spindle Manipulation) to +21.7% (Imagery 2), with significant within-experiment comparisons only for Imagery 2 (*p* = 0.014) and Spindle Manipulation (*p* < 0.001; S2 Table).

Both patterns combine in AF-Ratio: the drop from the stable to the unstable state was largest in the two experiments with the strongest AFiso_max_ reduction (Long COVID −48.7 pp, Spindle Procedure −47.8 pp) and smallest in Spindle Manipulation (−35.9 pp), but pronounced in all six experiments (all *p* < 0.001; S2 Table).

#### Joint model

With each factor adjusted for the other four, the association of muscle group persisted for AF_max_, AFiso_max_, AFosc and AF-Ratio (all *p* ≤ 0.017); for AF-Ratio and AFiso_max_ it was again carried by the three pectoralis major muscles (without them: *p* = 0.458 and *p* = 0.062). Of the remaining effects revealed in the single-factor models, only tester on AFosc (*p* < 0.001) and experiment on AFiso_max_ (*p* = 0.005) persisted; in addition, age became associated with AFiso_max_ and AF-Ratio (*p* = 0.035 and *p* = 0.039)—likewise disappearing without the pectoralis major. Because the factors are entangled, the joint model cannot assign an effect uniquely to one of them (S1 Table; Table 2).

For AF-Ratio, no factor remained associated with the difference between muscle states once the other factors were adjusted for and the pectoralis major muscles excluded (all *p* ≥ 0.062). The differences between subgroups were confined to the unstable state: across the 13 subgroups, AF-Ratio in the stable state stayed between 98.5 and 100.0%, whereas in the unstable state it ranged from 51.0 to 82.2% (51.0–62.6% excluding the pectoralis major muscles; S2 Table).

### Sensitivity analyses

Omitting each experiment in turn (82 to 102 muscles), excluding both spindle experiments, the imagery and odor experiments or the pectoralis major muscles, analyzing each tester separately, or restricting the analysis to the two experiments with blinded testers left the comparison between muscle states essentially unchanged (S3 Table). AF-Ratio was unaffected throughout, with MD between −41.6 and −46.0 pp (*d_z_* −2.49 to −2.90, all *p* < 0.001), as were AFiso_max_ (−100.0 to −57.7 N, *d_z_* −1.53 to −2.35), AFosc (+39.6 to 48.4 N, *d_z_* 1.18 to 1.44) and AFosc/AFiso_max_ (+1.1 to 1.4, *d_z_* 0.96 to 1.43; all *p* < 0.001). Only AF_max_ varied noticeably (+7.5 to 29.0 N, *d_z_* 0.24 to 0.85) and was the only parameter that approached the significance threshold (largest *p* = 0.048, after exclusion of the imagery and odor experiments); this was the only comparison that did not reach significance when considering the relative change between muscle states (+2.0%, *p* = 0.283).

Restricted to the two experiments in which the testers were blinded to the experimental condition (Imagery 2, Odors; 31 muscles of 21 participants), all five parameters still differed between muscle states (all *p* < 0.001, |*d_z_|* ranged from 0.85 (AF_max_) to 2.61 (AF-Ratio); S3 Table).

The association of muscle group with the difference in AF-Ratio rested on the three pectoralis major muscles: omitting one of them at a time shifted the unadjusted omnibus *p* from 0.003 to 0.013, 0.002 and 0.082, and excluding the group removed the association (*p* = 0.265).

### Illustrative single cases

Further AF data were collected in the context of imagery experiments and therapeutic practice (S1 Dataset): (1) a female student under current distress (exam pressure, 27 yrs., 174 cm, 69 kg; excluded from the imagery experiment), (2) a male patient with an acute ankle inflammation (64 yrs., 187 cm, 87 kg), (3) a male patient with a subluxation of the 5^th^ rib (65 yrs., 185 cm, 86 kg) and (4) a female participant (33 yrs., 167 cm, 56 kg) with a particularly pronounced reaction to negative food imagery. AF of elbow or hip flexors was measured analogously to the experiments described above (3 trials per state, except 2 unstable trials in Case 1; pre- and post-intervention on the same day within ∼60 min). Tester B performed measurements in Cases 1–3, tester A in Case 4. In Cases 1–3, the muscles were unstable in the entry state and regained stability after specific interventions: (1) positive imagery, (2) individualized pulsed electromagnetic field (PEMF) therapy to the ankle, (3) manual mobilization and PEMF to the rib. In Cases 2 and 3 the selection of treatment was guided by AF assessment [46]. Case 4 followed the imagery design with a stable state at baseline and instability induced by negative imagery.

Fig 6 illustrates representative signals of two AF trials per case, recorded from the identical muscle in both muscle states. The force increase was similar across stable and unstable trials, confirming the reproducibility of force application. Mean values of all trials per case are given in Fig 7.

**Fig 6.**
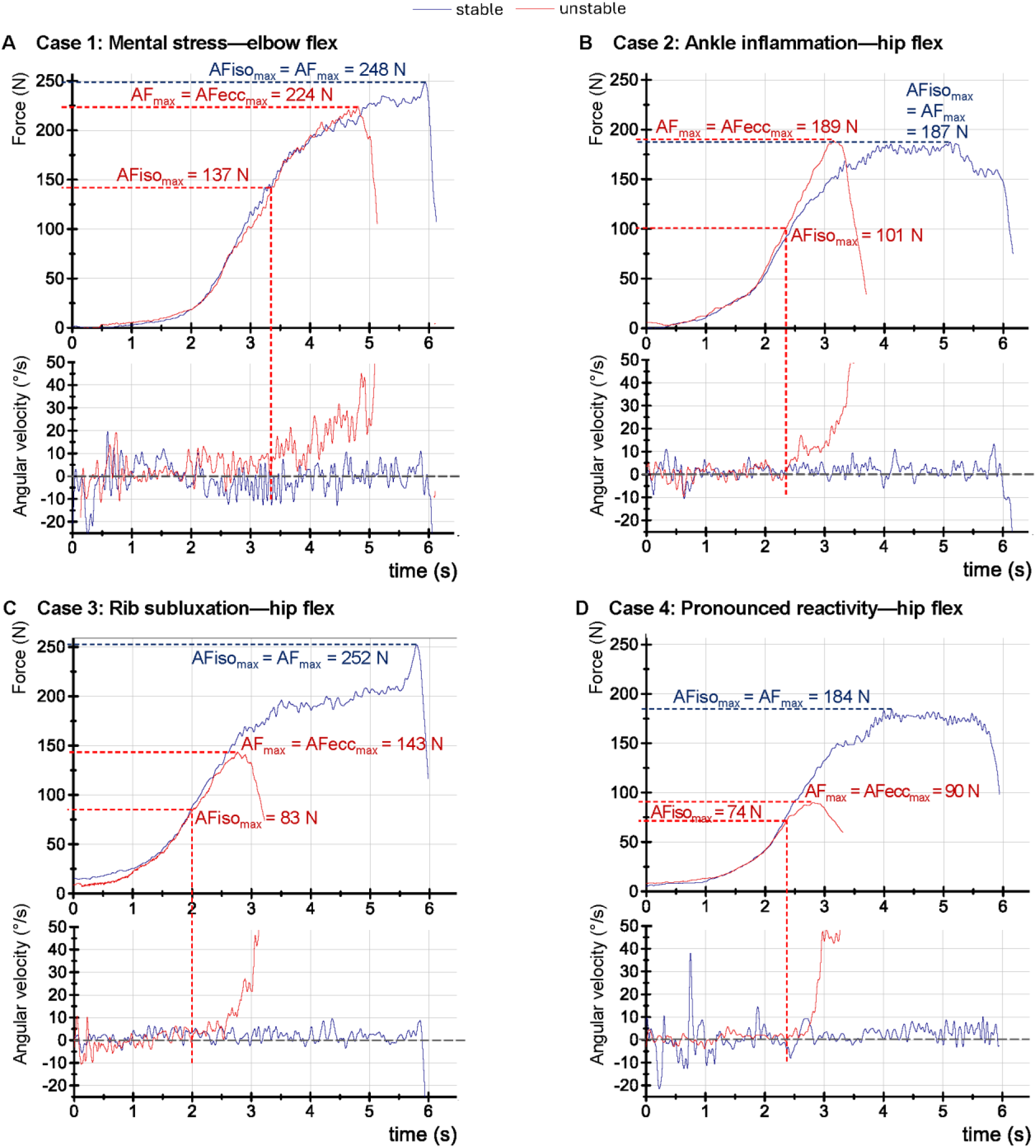
Single cases—representative signals of AF measurements. Displayed are the force (N) and angular velocity (°/s) signals of two AF trials per case, recorded from the identical muscle in the stable (blue) and the unstable (red) state. **(A)** Case 1—elbow flexors of a female student with current stress (unstable) vs. positive imagery (stable). **(B)** Case 2—hip flexors of a male patient with acute ankle inflammation at entry state (unstable) vs. post-treatment (stable) after pulsed electromagnetic field (PEMF) therapy. **(C)** Case 3—hip flexors of a male patient with rib subluxation at entry state (unstable) vs. post-treatment (stable) after manual mobilization and PEMF. **(D)** Case 4—hip flexors of a female participant during negative (unstable) vs. positive (stable) food imagery showing a particularly pronounced reaction to negative imagery. Marked are the max. isometric Adaptive Force (AFiso_max_) and peak AF (AF_max_). In case AF_max_ was reached during muscle lengthening, the term AFecc_max_ was used indicating eccentric muscle action.

**Fig 7.**
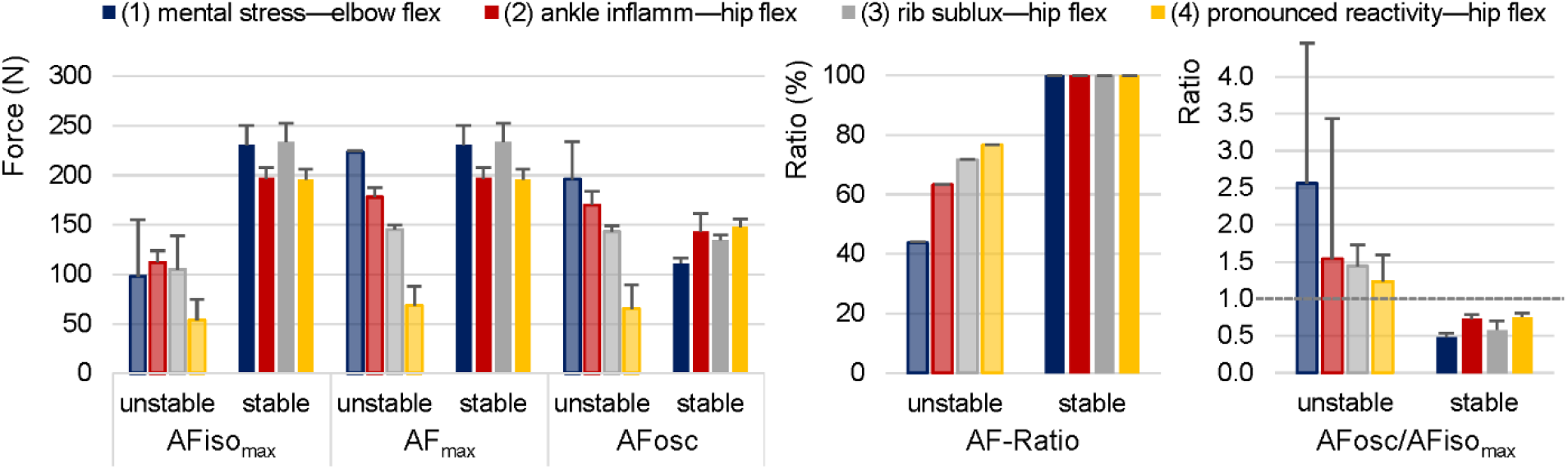
AF parameters of the four single cases. Displayed are the arithmetic means and standard deviations (error bars) of the trials per case in the unstable and stable state for the AF parameters AFiso_max_, AF_max_, AFosc (all in N), AF-Ratio (%) and AFosc/AFiso_max_; the dashed line in AFosc/AFiso_max_ marks the value of 1, above which oscillations arose after the break point. Case 1: mental stress (elbow flexors); Case 2: acute ankle inflammation (hip flexors); Case 3: rib subluxation (hip flexors); Case 4: pronounced reactivity to unpleasant imagery (hip flexors). AFiso_max_, max. isometric Adaptive Force; AF_max,_ peak AF; AFosc, AF at onset of oscillations; AF-Ratio, AFiso_max_/AF_max_ × 100.

In Cases 1 and 2, the pattern was consistent with the overall group: AF_max_ was at a high level for both states, while AFiso_max_ was clearly reduced for unstable muscles, resulting in averaged AF-Ratios of 44% and 63%, respectively (Figs 6A, B and 7). Oscillations arose after the break point in unstable and during the isometric phase in stable trials.

In Case 3, AF_max_ was markedly lower for unstable than stable muscles (on average 145 vs. 233 N; Figs 6C and 7), suggesting that the nociceptive stimulus of the rib subluxation limited force generation even during muscle lengthening. After treatment, AF_max_ recovered to a high level under stable conditions (= AFiso_max_). AF-Ratio in unstable state still averaged 72% (due to the particularly low AFiso_max_ of ∼105 N) and 100% for stable muscles, reflecting a similar pattern to the overall group despite the low AF_max_ in unstable trials. Oscillations arose only shortly before the peak force in unstable trials (AFosc/AF_max_ = 98%), while they occurred at 58% of AF_max_ in stable trials (Figs 6C and 7).

Case 4 showed the most pronounced reaction, mirrored not only by the holding capacity (AFiso_max_), but also by the peak force (Figs 6D and 7): during negative imagery, AF_max_ reached only 35% of AF_max_ in the stable state (∼68 vs. 196 N), with AFiso_max_ even lower (∼54 N). Although AF-Ratio (∼77%) was comparable to the overall group, it does not fully reflect the extent of the impairment in this case, as both AFiso_max_ and AF_max_ were dramatically reduced. A similarly pronounced reduction of AF_max_ in the unstable state was also observed in the elbow flexor of this participant (AF_max_ unstable vs. stable: ∼33%; not shown in Fig 7), indicating a generalized rather than muscle-specific response.

## Discussion

This study synthesized AF data from six studies comprising 788 trials of 112 muscles in 68 participants, with each muscle measured in both stable and unstable state. The most important findings are that (1) stable and unstable muscles can be clearly distinguished by the AF parameters, (2) the investigated factors (muscle group, tester, participant sex, age, experiment) modified the magnitude of the difference, and for AF-Ratio none of them persisted after adjustment for the others and exclusion of the pectoralis major, and (3) muscles can switch instantaneously between stability and instability in response to various stimuli. These findings on muscular adaptation are discussed below emphasizing the high potential for practical implications.

### Characterization of muscle stability: stable vs. unstable

The findings indicate that stable and unstable muscles can be clearly distinguished. Two basic key characteristics emerged:

1. Stable muscles are able to reach their full force capacity under isometric conditions during adaptation to an increasing external load (AF-Ratio: ∼99%), whereas unstable muscles yield at submaximal intensities (AF-Ratio: ∼56%). Notably, unstable muscles still reach—and even slightly exceed—their peak force of the stable state (AF_max_ +7.7%), but only during the subsequent eccentric phase. This suggests unstable muscles are not weak; they fail to access their full force capacity under static conditions, but retain the ability to generate high peak forces during muscle lengthening—a characterization of muscular instability (exceptions addressed below). This is further supported by two of the underlying studies and a more recent study, which additionally assessed MVIC: under stable conditions the holding capacity matched MVIC, whereas it fell to ∼55–67% of MVIC when the muscle was unstable [40,41]; in football players, AF_max_ even exceeded MVIC by 16% in unstable muscles while AFiso_max_ was 52% lower [47]. In that study, MVIC did not differ between the stability categories (*p* = 0.310, ω² = 0.00), whereas AF-Ratio accounted for 88% of the variance (ω² = 0.80)—indicating that the adaptive holding capacity constitutes a construct distinct from maximal strength [47]. These characterizations of stable and unstable muscles likely reflect differences in the adequacy of adaptive neuromuscular control—an interpretation to be developed further in the following sections. AF-Ratio best discriminates between the two states and proved robust across the investigated conditions: no factor remained associated with the difference between the states once the others were adjusted for and the pectoralis major was excluded (all *p* ≥ 0.062).
2. Stable muscles exhibit distinct mechanical oscillations already in the isometric phase at submaximal intensities (74% of AF_max_; in 474 of 479 trials the onset preceded the break point), whereas in unstable muscles no oscillatory onset was detectable in 29% of trials, and where it occurred, it was shifted close to the peak force (88% of AF_max_) and arose in the eccentric phase in 192 of 218 trials. Previous research has demonstrated that two coupled neuromuscular systems develop synchronized oscillations during isometric interaction at submaximal intensities [53–55]. The occurrence of oscillations could be a fundamental factor in creating muscle stability in a dynamic oscillating equilibrium as in the AF assessment (physiological basis considered below). When an interfering stimulus is present, the development of synchronized oscillations appears limited and largely fails under isometric conditions. Since the stimulus acts on the participant, who must adapt to the increasing force, the absent or delayed oscillatory onset likely originates on the participant’s side of the interaction. The onset of oscillations could be a sign of—or even a prerequisite for—muscular stability.

Table 3 summarizes the key characteristics of muscle stability compared to the conventional pushing strength.

**Table 3.**
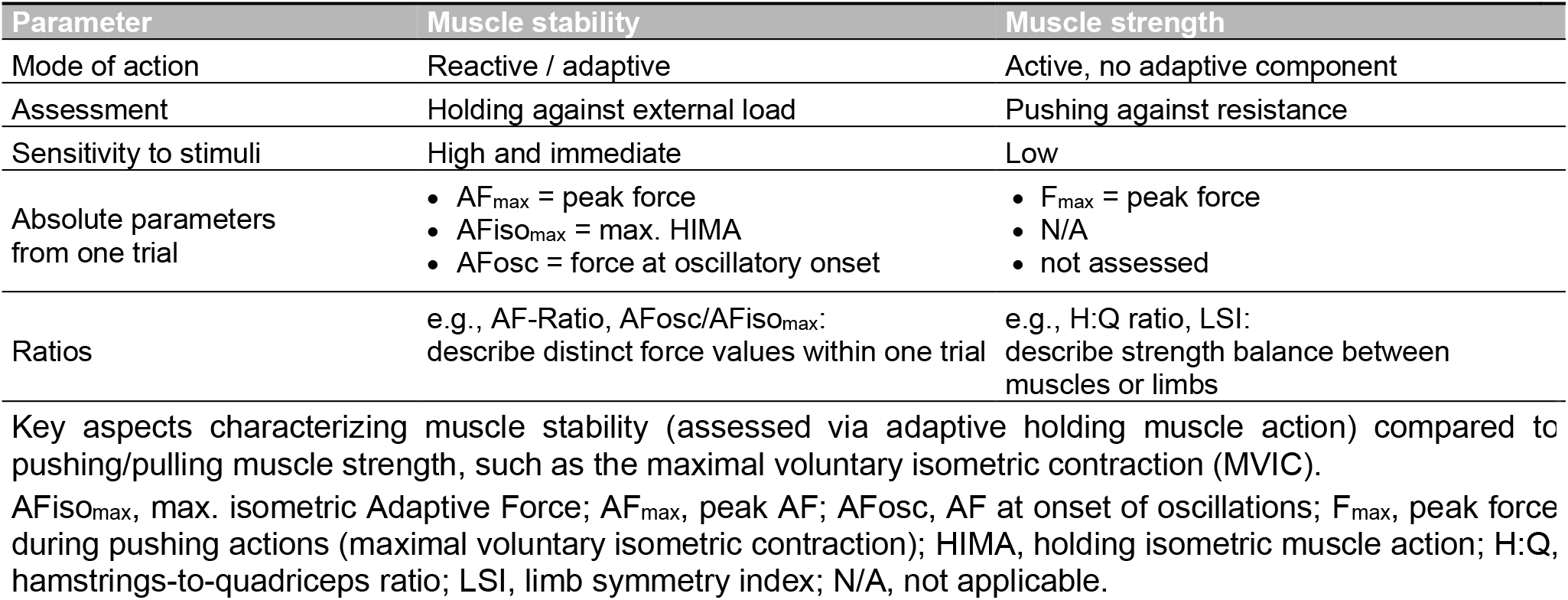
Conceptual comparison of muscle stability vs. muscle strength.

| **Parameter** | **Muscle stability** | **Muscle strength** |
|---|---|---|
| Mode of action | Reactive / adaptive | Active, no adaptive component |
| Assessment | Holding against external load | Pushing against resistance |
| Sensitivity to stimuli | High and immediate | Low |
| Absolute parameters from one trial | • AF<sub>max</sub> = peak force<br>• AFiso<sub>max</sub> = max. HIMA<br>• AFosc = force at oscillatory onset | • F<sub>max</sub> = peak force<br>• N/A<br>• not assessed |
| Ratios | e.g., AF-Ratio, AFosc/AFiso<sub>max</sub>:<br>describe distinct force values within one trial | e.g., H:Q ratio, LSI:<br>describe strength balance between muscles or limbs |

AFiso<sub>max</sub>, max. isometric Adaptive Force; AF<sub>max</sub>, peak AF; AFosc, AF at onset of oscillations; F<sub>max</sub>, peak force during pushing actions (maximal voluntary isometric contraction); HIMA, holding isometric muscle action; H:Q, hamstrings-to-quadriceps ratio; LSI, limb symmetry index; N/A, not applicable.

### Factors influencing parameters of muscle stability

A distinction must be made between factors related to the test constellation (e.g., physical conditions of tester and participants, muscle group, positioning, force application) and those that affect muscle stability physiologically (i.e., disruptive or supportive stimuli). The latter are discussed in subsequent sections.

Regarding test-related factors, a few basic physical principles should be mentioned first: During manual muscle testing, the measured force is always generated by both parties simultaneously, reflecting their combined performance. The tester must adjust the applied force to the participant’s capabilities and the muscle’s biomechanical requirements (e.g., less force is usually required for female participants, small muscles, or muscles tested with a long lever compared to males, large muscles, or short lever tests). Consequently, absolute force values depend on the tester-participant constellation.

In the present analysis, the tester was not associated with the difference between the muscle states for AFiso_max_, AF-Ratio, and AFosc/AFiso_max_ in any model; for AF_max_, an association with the tester emerged in the single-factor model across all 112 muscles, which did not persist after exclusion of the pectoralis major muscles or after adjustment for the other factors (all *p* ≥ 0.059). However, the absolute force levels differed markedly between the testers: in the stable state, AF_max_ was 36.4% lower for the female than for the male tester, with a corresponding value for AFiso_max_ (−36.2%; S2 Table). Despite this, the relative loss of AFiso_max_ in the unstable state was identical for both testers (−39.9% vs. −39.6%). This is the expected consequence of each muscle serving as its own control. The lower absolute values of the female tester could be attributed to differences in physical strength—females generally produce less maximum force than males [56]—however, both testers achieved similar peak forces against a fixed resistance (∼280 N) [39]. More relevant is the non-balanced distribution of participant sex and physical condition across testers—the male tester assessed predominantly male athletic participants in the spindle experiments—a confounding that cannot be disentangled with only two testers (Cramér’s V ≥ 0.56 for both; S1 Table).

The participants’ sex showed a similar constellation to the testers. In the stable state, AF_max_ and AFiso_max_ were ∼38% lower in females than in males. Descriptively, the loss of holding capacity in the unstable state was more pronounced in females, both for AFiso_max_ (−44.2% vs. −34.1%) and consequently for AF-Ratio (−48.0 vs. −38.0 pp; S2 Table). This is reflected in the single-factor association of participant sex with AF-Ratio across all 112 muscles, which did not persist after exclusion of the three pectoralis major muscles—all from male participants and all with unusually small AF-Ratio differences—or after adjustment for the other factors (all *p* ≥ 0.119). Since participant sex is itself unevenly distributed across the experiments (Cramér’s V = 0.74; S1 Table), a sex-related tendency cannot be separated from the test constellation in the present data and would require a balanced sample.

AFosc behaved differently: its change was associated with the tester in all models (*p* ≤ 0.010), but not with participant sex. The tester effect originates in the stable state: in unstable muscles, oscillations arose at nearly the same relative level irrespective of tester (∼91% of AF_max_), whereas in stable muscles their onset differed (male tester 79.8% vs. female tester 69.5% of AF_max_), resulting in a clearly different relative increase of AFosc (20% vs. 50%).

The different muscle groups showed the same qualitative pattern in the stable and the unstable state, indicating that the characteristics of muscle stability are consistent across different muscles; this consistency was also found across five muscle groups in football players, four of them not assessed here (knee flexors/extensors, hip adductors/abductors) [47]. Quantitatively, however, the muscle group was the factor most consistently associated with the difference between the states: the relative change of AF_max_ was larger for hip than for elbow flexors (12% vs. 4%), and this persisted after exclusion of the pectoralis major and after adjustment for the other factors; the same applied to AFosc, although this association was confirmed by the bootstrap but not by the permutation test (S4 Table). For AFiso_max_ and AF-Ratio, by contrast, the associations did not persist after exclusion of the pectoralis major (*p* = 0.019 vs. 0.083 and 0.014 vs. 0.796); for AF-Ratio the bootstrap did not confirm the association even across all muscles (*p* = 0.356). The markedly higher AF-Ratio of the three pectoralis major muscles in the unstable state compared to elbow and hip flexors (82% vs. 53% and 57%) may partly reflect how the break point was identified rather than a genuine difference in holding capacity: the pectoralis major was tested with a long lever, so the limb shows larger excursions during muscle testing, which may have shifted the gyroscope-based detection of the break point towards higher force values; the algorithm used was developed for elbow and hip flexors and did not account for this. With n = 3, and given the consistency across the other muscle groups assessed here and in football players, this appears specific to the test of the pectoralis major rather than a general muscle-group effect, and should be addressed in future studies with larger samples.

Age was not associated with the difference between the states in any single-factor model, and the slope per year of age was negligible for all parameters (S2 Table). Associations for AFiso_max_ and AF-Ratio emerged only in the joint model across all muscles (*p* = 0.035 and 0.039, unadjusted) and were absent after exclusion of the pectoralis major; given the close link between age and experiment (η = 0.61; S1 Table), these isolated signals more likely reflect the collinearity of the design than an effect of age. With a predominantly young sample (33.9 ± 13.1 years, range 19–70), age effects remain an open question that requires samples covering older participants.

Considering the factor experiment, the absolute AF values were highest in both spindle experiments (S2 Table)—likely attributable to the tester-participant constellation described above. The experiment was associated with the relative change of AFiso_max_ in all models: the loss of holding capacity ranged from 31% to 50% across the six experiments. Each of the two distribution-free checks confirmed this association in one of the two samples—the permutation test across all muscles, the bootstrap after exclusion of the pectoralis major. For AF_max_, an association emerged in the single-factor model but was not confirmed by either check (S4 Table). Crucially, AF-Ratio showed no experiment-related association in any of the analyses, confirming that the pattern of muscle stability and instability is consistent regardless of the type of intervention used to provoke it. Of note, only in the long COVID cohort was AF_max_—although not significantly—on average lower in the unstable than in the stable state (Fig 4, S2 Table); in all other experiments, unstable muscles generated similar or higher AF_max_ values during the eccentric phase (significant only for Spindle Manipulation and Imagery 2; S2 Table). The absence of this pattern in long COVID patients may indicate that in this condition, the overall force capacity is limited—not only the holding capacity. After treatment, muscle stability was restored immediately, with AFiso_max_ returning to the level of AF_max_ [45], suggesting that the limitation was reversible and functional rather than structural.

Beyond the factors examined, two extreme constellations emerged at the level of individual muscles. The pattern of markedly reduced peak force in the unstable vs. stable state—in addition to the reduction of AFiso_max_—is rare but clusters in identifiable subgroups. In 7 of 117 muscles (including single cases), the AF_max_ unstable/stable ratio fell below 70%. These included two female long COVID patients (3 muscles: 43.5–68.6%), the male participant with rib subluxation (Case 3: 62.3%)—in whom the affected rib was directly involved in the force transmission chain of the hip flexor test—one female participant from the imagery experiments (69.6%), and the female participant showing the most pronounced reduction (Case 4; elbow flex: 33.0%; hip flex: 35.0%). In both of the latter, instability was triggered purely by emotional imagery, suggesting particularly high individual reactivity. Together, these observations suggest that instability may, in some individuals, extend beyond the holding capacity to also affect peak force—potentially representing a further stage of the phenomenon that can arise from severe systemic conditions, local mechanical disruption during the test, particularly high individual reactivity, or a combination of these. Importantly, AF-Ratio itself remained within the expected range for unstable muscles even in these extreme cases, underscoring its independence from absolute force levels. At the other extreme, some individuals showed exceptionally low holding capacity (AF-Ratio <40% in 18 of 117 muscles from 14 participants; see Fig 5 and single cases), reaching values as low as 10%—meaning that when external forces act on the body, stabilization fails already at clearly submaximal levels, with relevance for daily life and sports activities where adaptation to external loads is required. Both phenomena—extremely low holding capacity and additional AF_max_ drop—showed no correlation (*r* = −0.05, *p* = 0.574; based on the 112 muscles of the main analysis), indicating that these represent two independent dimensions of muscle instability; both, however, appeared more often in female than male participants (AF-Ratio <40%: 10/14 female; AF_max_ unstable/stable <70%: 4/5 female).

In summary, differences in the absolute AF parameters are irrelevant as long as the same tester and setting are used for both muscle states, since each muscle serves as its own control. After exclusion of the pectoralis major muscles, no factor was associated with the difference of AF-Ratio in any model. This independence eliminates the need for reference values and makes AF-Ratio a robust and universally applicable marker, which also allows comparison between different studies.

### Neurophysiological considerations of muscle stability

The following considerations are presented as a working hypothesis: all statements on specific neural pathways are proposed mechanisms that require direct experimental investigation. They are necessarily simplified given the complexity of the control networks involved, but outline the key mechanisms that may explain the observed phenomena.

The adaptive holding function requires a continuous sensorimotor feedback loop involving proprioceptive afferents (muscle spindle cells, Golgi tendon organs, joint and skin receptors) and a complex network of central structures including the sensorimotor cortex, thalamus, cerebellum, inferior olivary nucleus, basal ganglia, and cingulate cortex [27,57–61]. A mixed feedforward–feedback control mechanism is assumed, in which the cerebellum—in cooperation with the inferior olive—acts as a forward controller, predicting the sensory consequences of motor commands and enabling temporally coordinated adjustments [27,57,62,63]. For optimal AF execution, the system must continuously compare the intended motor command (efference copy) with the actual sensory feedback (reafference) and correct any mismatch by adjusting motor output [27,64].

This control architecture may explain why the holding function is more demanding than purely force-generating actions: when pushing or pulling, the force target is self-generated and does not require continuous adaptation to a varying external input, placing relatively lower demands on sensorimotor control. During holding actions—and particularly during AF assessment with an increasing external load—the system must constantly update its motor output based on changing sensory input, placing higher demands on the cerebellar–olivary timing circuitry and the proprioceptive feedback loop [27,30,57].

Crucially, several of the structures involved in this sensorimotor control network also process emotional, olfactory, and nociceptive information. The amygdala modulates motor responses to threat perception via connections to supplementary motor areas [65]. The anterior cingulate cortex integrates executive motor functions with emotional processing [61,66]. The cerebellum is involved in both adaptive motor control [58,60] and olfactory processing [67,68]. Nociception directly affects motor output through spinal and supraspinal pathways [69]. The thalamus serves as a central integrative hub—far beyond a simple relay station—through which nearly all sensory, nociceptive, and limbic information must pass before reaching the cortex, while simultaneously receiving feedback from motor control structures including the cerebellum and basal ganglia [60,70–75]. This convergence of diverse inputs within a single central structure provides a plausible neuroanatomical basis for the observed effects of emotional, olfactory, proprioceptive, and nociceptive stimuli on adaptive motor control.

The instantaneous nature of the stability–instability switch is consistent with a reflexive mechanism operating through these fast-conducting pathways rather than through voluntary cognitive processes. The reproducibility of this response across the underlying studies—where adequate impairing stimuli consistently triggered instability and their removal restored stability—supports a stimulus-dependent reflexive mechanism. The long-loop latency of ∼80–100 ms through the cerebellar–cortical circuits [64] would be sufficient for such rapid modulation. It is hypothesized that disruptive stimuli interfere with the target–actual comparison of muscle length and tension, disrupting the forward model and causing the system to fail at maintaining the isometric condition above a certain force threshold, i.e., at readjusting length and tension adequately.

The gamma motor system may provide a specific mechanism for this selective impairment of holding capacity with preserved maximal force in response to disruptive stimuli. Under physiological conditions, alpha and gamma motor neurons are coactivated in a task-dependent manner, ensuring that muscle spindles remain sensitive during active contractions and continue to provide accurate length information [76]. This alpha–gamma coactivation is presumably calibrated by the cerebellum, which integrates efference copies of motor commands with spindle afference via spinocerebellar pathways [77]. The tonic gamma drive is substantially regulated by the reticulospinal tract [78]. Its origin in the pontomedullary reticular formation serves as a convergence point for modulating input from stress-related axes, the autonomic nervous system, and limbic structures [79,80], along with nociceptive afferents relayed via the spinoreticular pathway [78,81]. This pathway thus represents a physiologically plausible route through which psychovegetative states—including emotional distress, post-infectious dysregulation, and autonomic imbalance—as well as nociceptive input may alter gamma motor neuron excitability and, consequently, spindle sensitivity. A disturbance of this gamma-mediated calibration would specifically compromise the target–actual comparison of muscle length outlined above, leading to inadequate force adjustment and thus the observed instability during adaptive holding actions.

This proposed mechanism is supported by the spindle manipulation experiments [40,41], in which selective disruption of spindle afference at the peripheral level reproduced the characteristic instability pattern without impairing peak force capacity—indicating that disturbed spindle-level proprioception alone can be sufficient to produce the phenomenon. A central route converging on the same mechanism is indicated by the observation that emotional [42,43] and olfactory stimuli [44] as well as the long COVID condition (associated with autonomic dysregulation) [45] elicited comparable instability, though the specific neural pathways involved remain to be established. While these proposed underlying pathways require further investigation, the convergence of spindle-level, spinal, and supraspinal mechanisms within the reticulospinal–gamma axis offers a parsimonious framework for the diverse stimuli observed to affect muscle stability.

The control loop outlined above operates cyclically, and its central structures and the muscular output oscillate in similar frequency ranges: the olivocerebellar system, the deep cerebellar nuclei, and the thalamic motor nuclei all show rhythmic activity in overlapping ranges of ∼8–17 Hz, with the inferior olive regarded as a pacemaker of rhythmic cerebellar activity [62,82–84], while the mechanical micro-oscillations of muscle are stochastically distributed around 10 Hz (range ∼8–20 Hz) [85–89]. AFosc denotes the force at the onset of such oscillations; the criterion used to identify this onset is a conservative lower bound and does not resolve the frequency. Since the effective neural drive to the muscle replicates the synaptic input common to the motoneuron pool [90], the mechanical oscillations carry the central drive. Rhythmic motor output has accordingly been used as a window into central oscillatory control [86]; its central origin is further supported by its responsiveness to transcranial magnetic stimulation, its alteration after central nervous lesions and in Parkinson patients without manifest tremor [91–96]. Within this framework, oscillations arising at submaximal levels during the isometric phase in stable muscles would mirror adequately functioning central circuitry, whereas their absence, their delayed onset or their appearance only during yielding in unstable muscles would indicate a failure of this oscillatory control—an error in the sensorimotor loop rather than a reduced force-generating capacity.

The control architecture outlined in this section would also explain why the stability–instability switch responds to such diverse inputs, and why it does so immediately and reversibly—properties that carry implications for screening, prevention, diagnostics, and therapeutic measures.

### Relevance for musculoskeletal complaints and non-contact injuries

Given the high prevalence of musculoskeletal complaints and non-contact injuries in the general population and athletes with substantial social and economic burden [1–6], suitable diagnostic, preventive, and therapeutic approaches are urgently needed. Despite decades of research, the causes of musculoskeletal complaints and non-contact injuries remain insufficiently understood, with research largely confined to the description of risk factors. Maximum strength and neuromuscular imbalances have not proven to be reliable predictors [16–26]. The commonly cited concept of ‘overload’ does not provide a plausible explanation for why complaints often arise unilaterally despite symmetrical loading, why only a minority of athletes develop injuries under equal training conditions, or why well-trained athletes develop complaints or non-contact injuries at all with normal exertion [97]. Load is certainly a factor, but intrinsic factors affecting load capacity must be involved: where this capacity is reduced, even loads that are usually tolerated may suffice to produce complaints or injuries.

A key observation is that non-contact injuries and musculoskeletal complaints predominantly occur during activities requiring active deceleration and dynamic stabilization [7–16]—situations in which muscles must hold or brake against external forces. Conventional strength assessments using pushing actions without reactive components do not capture this adaptive holding function. The present findings suggest that muscle instability—a reflexive, stimulus-dependent reduction of the holding capacity—could represent a missing factor of high relevance in this context. Under load, joint stability depends on the muscles as active stabilizers, which thereby protect the passive structures [98]. When the holding capacity is substantially reduced, the affected muscle may become the weakest link in the kinetic chain, yielding prematurely, so that joints and connective tissue structures are exposed to loads they are not designed to withstand—plausibly resulting in complaints and non-contact injuries. Accordingly, intact muscle stability may be a precondition for protecting these structures under load and thus for preventing and resolving such injuries and complaints. We propose the term functional instability syndrome (FIS) to describe this mechanism. It is hypothesized that muscle instability represents a necessary—though not necessarily sufficient—condition for the development of musculoskeletal complaints and non-contact injuries. FIS remains a hypothetical construct that requires verification through prospective studies.

Several aspects of the present data, however, support this hypothesis. First, the holding capacity reacted sensitively and instantly to various disruptive stimuli, with clear reductions of on average 43.6 pp (AF-Ratio 99.4% to 55.8%). The holding function can thus be compromised while maximal strength remains preserved [40,41,47,98,99]—which may explain why strength-related parameters show no consistent association with complaints and injuries [16–26]. Second, the response was reversible: stability was restored immediately after removal of the impairing stimulus, presentation of a supportive stimulus, or after targeted therapy. This suggests a predominantly functional nature, which could explain why musculoskeletal complaints frequently occur without a structural correlate [100,101]. Conversely, preserved muscle stability might explain why structural findings are frequently present in asymptomatic individuals [102,103]. Third, the high variation of AF-Ratio in the unstable state (range 10.3–91.4%, SD 16.0 pp vs. 1.8 pp in the stable state) reflects individual differences in regulatory capacity and sensitivity in reaction to disruptive stimuli, with some individuals reacting particularly strongly, which might indicate a higher susceptibility to complaints and injuries. Beyond such individual predispositions, the instantaneous nature of the stability–instability switch further suggests that situational factors—such as acute emotional stress, pain, or sudden disruptive sensory input—may transiently compromise stability even in otherwise unaffected individuals. This carries particular relevance for sport, where such acute state shifts in critical moments could contribute to sudden non-contact injuries. Fourth, the impairing effect of proprioceptive irritation and nociception on muscle stability—investigated in [37,40,41,45] and illustrated by the presented case examples—may explain the reported high re-injury rates, predominantly non-contact in nature [18,20,104–106], and might also be relevant to the high risk of post-traumatic osteoarthritis [107–109]. Structural damage such as hamstring strain [18], ACL rupture and reconstruction [20,104], or concussion [106,110,111] can cause residual proprioceptive irritation and nociception even after months, affecting neuromuscular control; this could in turn affect the holding capacity beyond the directly affected area and thereby increase vulnerability to injury under load. The link between stability deficits and musculoskeletal complaints was found in football players fully participating in training and matches, that is, at a stage at which complaints had not led to time loss [99]. Moreover, in long COVID patients the holding capacity was restored in the course of treatment, while the intensity of most reported symptoms, including muscle pain, decreased significantly from the long COVID state to the end of the treatment series [45]. Both changes occurred in the same patients, although their association was not formally tested. Fifth, the connection between psychosocial stress and musculoskeletal complaints is well-known, yet the underlying mechanisms are not resolved [112–116]—here too, a reduced holding capacity caused by emotional stressors could provide an explanation. Further investigations are justified and needed.

The evidence to date suggests that the assessment of muscle stability in the sense of AF—being highly sensitive and responding instantaneously to supportive or impairing stimuli—holds potential as a tool for screening, prevention, targeted diagnostics, and for deriving individualized therapy from the AF response (AF-guided therapy). Manual muscle testing has long been used in clinical practice, albeit based on the examiner’s subjective evaluation and largely without standardization; a handheld device recording force and position simultaneously, combined with a standardized test protocol, now allows this manual test to be objectively quantified and investigated systematically.

Although AF-Ratio is a continuous measure, the stable and unstable states separated almost completely in the present data. Clinical use requires a continuous variable to be translated into a dichotomous outcome [22]—a step that here does not depend on an arbitrarily set cut-off, since the two states appear as distinct qualities rather than as a gradient. In football players, AF-Ratio discriminated those with and without musculoskeletal complaints (AUC 0.93) and, in preliminary follow-up, those with subsequent complaints or injuries (AUC 0.86, n = 13), whereas MVIC, the hamstring-to-quadriceps ratio and AF_max_ did not discriminate in the hypothesized direction (AUC ≤ 0.44 cross-sectionally, AUC ≤ 0.38 at follow-up) [99]. Since AF and its standardized assessment were introduced by the present research group, the available evidence necessarily derives from this group and from a measurement system developed by it; the corresponding patent is held by the University of Potsdam (see Competing interests). Independent replication by other research groups is therefore an essential next step.

If muscle instability is a necessary condition for the development of musculoskeletal complaints and non-contact injuries, its restoration would be a necessary step in resolving or preventing them. Within the FIS framework, the AF-guided treatment approach—in which interventions are selected based on their ability to restore muscle stability—would open considerable potential for individualized, targeted, cause-oriented treatment. Its procedure has been described previously in long COVID patients [45,46] and was applied in Cases 2 and 3 presented here. Further clinical investigation is warranted.

Regarding prevention, two aspects should be addressed: First, the holding capacity is affected early—deficits occur even in asymptomatic participants—which would make the assessment a candidate for screening before complaints or injuries arise. Second, the reflexive nature of the instability response implies that muscle stability may not be trainable in the conventional sense. However, two complementary strategies emerge: (1) identification and elimination of individual disruptive stimuli leading to instability through AF-guided diagnostics and cause-related individualized therapy, and (2) optimization of the adaptive holding and braking function through targeted training programs. For example, a preventive effect on hamstring strain injuries was suggested—although still debated—for the Nordic hamstring exercise [117–121], which includes adaptive components and may improve AF. Cases with limited effectiveness could indicate the presence of an underlying cause that affects the holding capacity and cannot be resolved by exercise. Further research should address these aspects.

In summary, the concept of muscle stability shifts the focus from load to individual capacity: what matters is not primarily strength, but whether the neuromuscular system can maintain adequate control under external loads. Given the limited value of strength screening for injuries and complaints, the AF assessment offers a novel framework addressing a neuromuscular function challenged during injury-prone movements. The concept provides plausible explanations for unresolved questions in sports science and medicine and opens up possibilities for screening, targeted diagnostics, and AF-guided therapy.

### Limitations and methodological aspects

Limitations of the underlying studies are discussed in the respective articles. The present analysis used pre-sorted data from six studies to identify common characteristics of stable and unstable muscles. This approach does not allow conclusions about stimulus-specific responses (reported in the original studies). The present sample, though adequate for identifying fundamental characteristics, is limited in size and diversity. Generalizability to other populations (e.g., different age groups, sports, pathologies) requires verification.

The classification into stable and unstable states was based on the testers’ clinical ratings, made without knowledge of the recorded signals (see Methods). Rating and AF parameters necessarily refer to the same mechanical event—the yielding of the limb—since the measurement serves to quantify what the tester perceives; their agreement therefore expresses the degree of objectification. As muscle stability in the sense of AF is a newly introduced construct, no established reference standard exists against which the parameter could be validated. Three lines of evidence are independent of the testers’ ratings. First, the condition-based classification rests on the experimental manipulation alone and reproduced all five comparisons—known-groups validity against an experimentally defined state. Second, the threshold was defined in the odor study [44] and thus applied prospectively to the four experiments published later (643 trials; agreement 97.4%, κ = 0.95). Third, the parameter discriminates where the established standard does not: in football players, MVIC and the hamstring-to-quadriceps ratio did not differ between the stability categories, whereas AF-Ratio did [47]—evidence of discriminant validity, that is, of a construct distinct from maximal strength. What remains to be established is predictive validity against clinical outcomes; a first step is reported for the same cohort [99]. The threshold of AF-Ratio ≥98% for stable state served here only to verify the agreement between rating and objective data; validated cut-off values for clinical decisions must be established in independent samples.

The effects of tester and participant characteristics on AF values could not be fully disentangled due to the non-balanced tester-participant constellation, as discussed above. Although tester strength could limit the test, the tester can use the entire body while the participant’s muscle is largely isolated, and the strength demands are moderate given the pronounced instability of impaired muscles—so the tester’s maximal strength is subordinate to the test outcome. A prerequisite for tester independence, the reproducibility of the applied force profile against fixed resistance, has been shown within and between the two testers (ICC(3,1) = 0.989 [39]); AF-Ratio was moreover not associated with the tester in any model, with virtually identical relative losses of AFiso_max_ (−39.9% vs. −39.6%) despite differences in sex, age, and physical condition. Reproducibility across further testers, and of the ratings themselves, is nevertheless not established. One consideration applies to conventional inter-rater designs: muscle stability can change instantly in response to stimuli, so disagreement between two successive raters may not be attributable to the raters alone. An inter-rater study accounting for this is planned.

Because of the small number of pectoralis major muscles (n = 3), all factor analyses were additionally run without this muscle group, revealing that the association of muscle group and participant sex with the difference in AF-Ratio rested entirely on these three muscles. As discussed above, the deviating pattern of pectoralis major is more likely attributable to data extraction adopted unchanged from the flexor muscles than to muscle-specific differences; the break point detection via angular velocity requires refined data processing for muscles tested with a long lever. A first step in this direction was taken in the football study, which implemented a novel algorithm across several muscle groups [47,99].

Within the underlying experiments, the causal direction is established for the stimulus: impairing stimuli induced instability, and their removal or a supportive stimulus restored stability. What the present design cannot establish is the causal link to musculoskeletal complaints and non-contact injuries. Measures of diagnostic accuracy were not determined in the present analysis, which the design does not permit; for the football cohort they are reported elsewhere [99]. The four single cases are uncontrolled observations that were not part of the inferential analysis; they illustrate response patterns and cannot demonstrate therapeutic effectiveness. Prospective studies are needed to establish whether muscle instability—as assessed by AF—predicts the occurrence of musculoskeletal complaints or injuries; a preliminary follow-up revealed first promising hints [99]. Clinical studies are required to evaluate the effectiveness of AF-guided diagnostics and individualized therapy.

### Conclusion

Stable and unstable muscles can be clearly distinguished by AF parameters, most consistently by AF-Ratio. In the unstable state, the holding capacity dropped markedly while peak force was preserved or even increased during the subsequent eccentric phase: in most cases, unstable muscles are not weak—they fail to access their full force capacity in the isometric position, so the deficit concerns neuromuscular control rather than strength. For AF-Ratio of hip and elbow flexors, no factor was associated with the difference between the two muscle states. AF-Ratio is self-referenced to the muscle’s own peak force and therefore requires no normative reference values.

The holding capacity responded instantly to disruptive stimuli and was restored equally rapidly upon their removal, supporting a reflexive, stimulus-dependent mechanism. Building on this, we propose the functional instability syndrome (FIS) as a hypothetical mechanism linking muscle instability to the development of musculoskeletal complaints and non-contact injuries; the neurophysiological account outlined above is a working hypothesis. AF assessment offers considerable potential for screening, prevention, targeted diagnostics, and individualized AF-guided therapy; its clinical value must now be evaluated in prospective and clinical studies. Shifting the focus from strength to muscle stability and from load to individual capacity offers a plausible explanation for the hitherto insufficiently understood development of musculoskeletal complaints and non-contact injuries.

## Acknowledgments

We would like to thank all student and research assistants as well as the participants for their active support and participation in the underlying studies.

## Supporting Information

### S1 Dataset. Trial-level Adaptive Force data

Anonymized values of AF_max_, AFiso_max_ and AFosc for all 788 trials of 112 muscles from 68 participants, with muscle group, tester, participant sex, age, experiment, the tester-rated muscle state and the condition-based classification, together with a description of all variables.

### S1 File. Analysis script

Python script reproducing every number reported in this article from S1 Dataset.

### S1 Fig. Q–Q plots of the within-muscle differences between the two muscle states

Standardized within-muscle differences (unstable − stable) of the five AF parameters and of the log-transformed ratio AFosc/AFiso_max_ plotted against the theoretical quantiles of the normal distribution (n = 112 muscles). The line marks exact normality; Shapiro–Wilk W and *p* are given in each panel.

### S1 Table. Composition of the experiments and separability of the potential influencing factors

Distribution of the four categorical factors and of age across the six experiments (S1A Table), the pairwise association between the factors (S1B Table) and the variance inflation of each factor in the joint model of Table 2 (S1C Table). Counts refer to muscles (n = 112); age refers to participants (n = 68).

Cramér’s V measures how strongly two categorical factors determine one another, from 0 (independent) to 1 (one factor fully determined by the other); the bias-corrected form is given. The number of empty cells states how many factor combinations do not occur at all, so that the corresponding adjusted contrast of the joint model rests on no observations. For the continuous factor age the correlation ratio η is given, which is interpreted on the same scale.

The generalized variance inflation factor GVIF quantifies by how much the estimation of a term is inflated by the remaining terms of the joint model. Because GVIF grows with the number of degrees of freedom, terms are compared through GVIF^1/(2df)^; the resulting inflation is modest: the largest value, 1.70 for participant sex, corresponds to a standard error about 1.7 times that expected under independent factors. All factors remain below 2.24, corresponding to a variance inflation factor of 5, so the joint model is estimable, although the factors are strongly entangled.

df, degrees of freedom; GVIF, generalized variance inflation factor; M, mean; n, number; SD, standard deviation; η, correlation ratio.

### S2 Table. Comparison of the stable and the unstable muscle state within each subgroup

Muscle-level values (n = 112 muscles of 68 participants) for every subgroup of the four categorical factors, separately for each AF parameter (S2A–S2E Table), with age covered separately in S2F Table (continuous covariate, no subgroups). Given are the means ± standard deviations of the two muscle states, the mean within-muscle difference MD (unstable − stable; percentage points for AF-Ratio) with its 95% confidence interval, the *p*-value of the paired t-test on MD, and the effect size *d_z_* with its noncentral-t 95% confidence interval. For the three force parameters the relative change, calculated per individual muscle as (unstable − stable)/stable × 100 and averaged across the muscles of the subgroup, is additionally reported; this is the quantity compared between subgroups in Table 2 and Fig 4, for which the MD itself serves in the case of AF-Ratio and AFosc/AFiso_max_. The reference row “All muscles” repeats the overall values of Table 1; its *p*-value is unadjusted.

Two different tests are reported. The *p* of the columns refers to the comparison of the two muscle states within one subgroup and is Holm-adjusted within each parameter across the 13 tests of that table (3 muscle groups, 2 testers, 2 sexes, 6 experiments). All confidence intervals are unadjusted for multiplicity, so an interval excluding zero does not necessarily correspond to a *p*-value below 0.05 after Holm adjustment; this occurs in three subgroups of S2A Table (AF_max_). These within-subgroup tests describe how consistent the difference is across subgroups and are not intended as confirmatory evidence, which is provided by Table 1 for the difference itself and by Table 2 for its modification by the factors.

The *p* of the shaded factor rows refers to the comparison between the subgroups of that factor; it is taken unchanged from Table 2, where Holm is applied within each parameter across the five factors, and is accompanied by the value obtained after exclusion of the pectoralis major. Each adjustment covers the tests actually carried out for the respective question, so the two sets of *p*-values are not interchangeable.

Confidence intervals for *d_z_* are noncentral-t intervals rather than the cluster bootstrap used in Table 1, because subgroups comprise as few as three participants; both methods agree closely where both apply (AFiso_max_, all muscles: noncentral t [−2.20; −1.58] vs. cluster bootstrap [−2.34; −1.61]). Estimates for the pectoralis major rest on three muscles of three participants and are reported for completeness only.

In S2F Table (age), b denotes the slope per year of age from a mixed model with a random intercept for participant, fitted to the same quantity that is compared between subgroups in Table 2 and Fig 4, with *p* Holm-adjusted across the five factors as in Table 2.

AFisomax, maximal isometric Adaptive Force; AFmax, peak Adaptive Force; AFosc, Adaptive Force at the onset of oscillations; AF-Ratio, AFisomax/AFmax × 100; b, regression slope; CI, confidence interval; *dz*, effect size for paired samples; M, mean; MD, mean difference; N, Newton; n, number; *p*, *p*-value; pp, percentage points; SD, standard deviation.

### S3 Table. Sensitivity analyses: the comparison between muscle states on restricted samples

The main comparison of the difference between muscle states (unstable – stable) for each AF parameter (S3A to S3E) was repeated after omitting each experiment in turn, after omitting subgroups defined by content, for each tester separately, and restricted to the two experiments in which the testers were blinded to the experimental condition. Given are the number of muscles and participants of each restricted sample, the mean within-muscle difference (MD, unstable minus stable), the effect size d_z_ and the *p*-value of the paired t-test.

*P*-values are unadjusted: these analyses probe the robustness of the pooled result and do not test new hypotheses. The long COVID cohort is the only experiment with diseased participants; its omission is listed under “One experiment omitted”. The full sample is given as reference in the first row.

AFisomax, maximal isometric Adaptive Force; AFmax, peak Adaptive Force; AFosc, Adaptive Force at the onset of oscillations; AF-Ratio, AFisomax/AFmax × 100; *dz*, effect size for paired samples; MD, mean difference; N, Newton; n, number; *p*, *p*-value; pp, percentage points; w/o, without.

### S4 Table. Distribution-free checks of the factor analyses

Two distribution-free counterparts of Table 2, both covering the single-factor models. Each factor is reported for all 112 muscles and after exclusion of the three pectoralis major muscles. The outcome is the relative change per muscle for the three force parameters and the difference for AF-Ratio and AFosc/AFiso_max_. All p-values are Holm-adjusted within each parameter across the five factors as in Table 2. Values below 0.05 are set in bold.

A. Permutation test. It makes no assumption about the distribution of the outcome. Under the null hypothesis that a factor is unrelated to the change between the muscle states, the participants are exchangeable. The complete label vector of a participant was therefore permuted as a block among the participants contributing the same number of muscles, which preserves the cluster sizes and the within-participant pattern of the muscle group. The test statistic is the Wald χ² of the same omnibus contrast as in Table 2, from the same random-intercept model; the variance components do not depend on the factor labels under the null and were estimated once per parameter from the intercept-only model, so that the observed and the permuted statistics are formed in exactly the same way. Given is *p* (χ²), with *p* = (1 + number of permutations with χ² ≥ observed) / 20,001 (Monte Carlo error ± 0.0015 at *p* = 0.05).
B. Wild cluster bootstrap. It addresses unequal residual variances rather than non-normality. The same omnibus contrast was fitted by ordinary least squares with the covariance clustered on participant, which is valid under both heteroscedasticity and within-participant dependence. Since cluster-robust Wald tests reject too often when a factor is identified from few clusters per level—the six experiments comprise 9 to 17 participants—the reference distribution was obtained by the wild cluster bootstrap with the null imposed and Rademacher weights: y* = fitted value + w × residual of the restricted model, with w drawn from {−1, +1} once per participant. Given is *p* = (1 + number of resamples with F* ≥ observed F) / 10,000 (Monte Carlo error ± 0.002 at *p* = 0.05).

AFisomax, maximal isometric Adaptive Force; AFmax, peak Adaptive Force; AFosc, Adaptive Force at the onset of oscillations; AF-Ratio, AFisomax/AFmax × 100; df, degrees of freedom of the omnibus contrast; p, p-value; w/o, without; χ², Wald chi-squared statistic.

