## Supplementary figures and images for "From strength to stability: muscular holding capacity characterized by Adaptive Force and its potential relevance for musculoskeletal complaints and non-contact injuries"

### S1 Fig

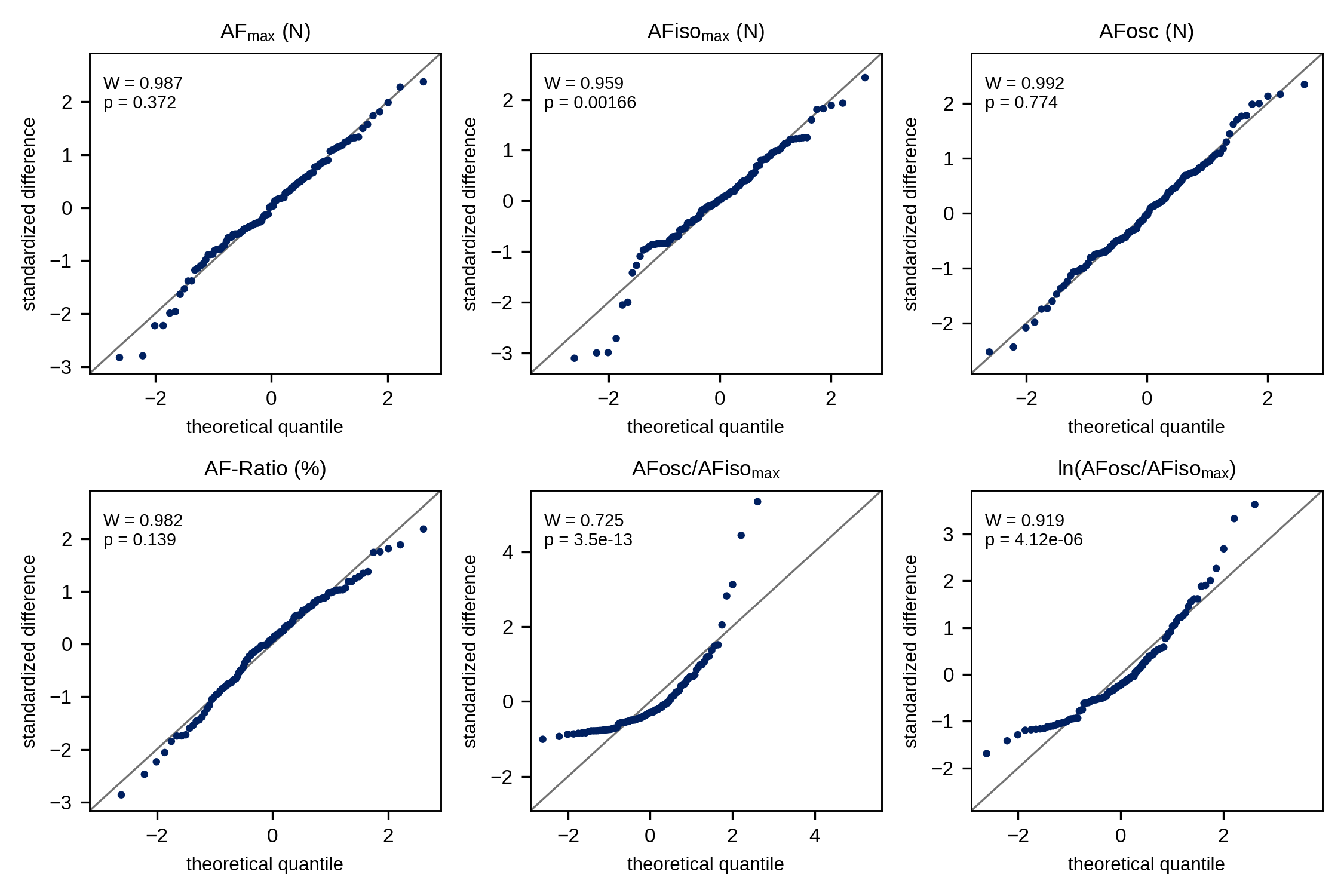
