## Supplementary material for "From strength to stability: muscular holding capacity characterized by Adaptive Force and its potential relevance for musculoskeletal complaints and non-contact injuries": S1 Table

Schaefer LV, Bittmann FN

#### S1 Table. Composition of the experiments and separability of the potential influencing factors.

Distribution of the four categorical factors and of age across the six experiments (S1A Table), the pairwise association between the factors (S1B Table) and the variance inflation of each factor in the joint model of Table 2 (S1C Table). Counts refer to muscles (n = 112); age refers to participants (n = 68).

Cramér's V measures how strongly two categorical factors determine one another, from 0 (independent) to 1 (one factor fully determined by the other); the bias-corrected form is given. The number of empty cells states how many factor combinations do not occur at all, so that the corresponding adjusted contrast of the joint model rests on no observations. For the continuous factor age the correlation ratio η is given, which is interpreted on the same scale.

The generalized variance inflation factor GVIF quantifies by how much the estimation of a term is inflated by the remaining terms of the joint model. Because GVIF grows with the number of degrees of freedom, terms are compared through $GVIF^{1/(2df)}$; the resulting inflation is modest: the largest value, 1.70 for participant sex, corresponds to a standard error about 1.7 times that expected under independent factors. All factors remain below 2.24, corresponding to a variance inflation factor of 5, so the joint model is estimable, although the factors are strongly entangled.

df, degrees of freedom; GVIF, generalized variance inflation factor; M, mean; n, number; SD, standard deviation; η, correlation ratio.

### S1A Table. Composition of the experiments.

| Experiment | n muscles | n partici-pants | Age (years) M ± SD (range) | Sex f / m | Tester A / B | Muscle group elbow / hip / pect |
|---|---|---|---|---|---|---|
| Imagery 1 | 10 | 9 | 38.6 ± 18.9 (22–70) | 6 / 4 | 3 / 7 | 1 / 6 / 3 |
| Imagery 2 | 21 | 12 | 25.0 ± 3.6 (20–31) | 21 / 0 | 0 / 21 | 10 / 11 / 0 |
| Odors | 10 | 9 | 34.1 ± 11.9 (24–57) | 6 / 4 | 4 / 6 | 2 / 8 / 0 |
| Spindle Procedure | 18 | 11 | 31.5 ± 8.6 (22–52) | 4 / 14 | 18 / 0 | 18 / 0 / 0 |
| Spindle Manipulation | 23 | 13 | 26.4 ± 4.3 (19–34) | 0 / 23 | 23 / 0 | 11 / 12 / 0 |
| Long COVID | 30 | 17 | 45.2 ± 13.7 (21–63) | 26 / 4 | 12 / 18 | 14 / 16 / 0 |
| **Total** | **112** | **68†** | **33.9 ± 13.1 (19–70)** | **63 / 49** | **60 / 52** | **56 / 53 / 3** |

†*Three participants took part in two experiments (Imagery 1 and Odors), contributing one muscle to each; the column therefore sums to 71 while 68 individuals were measured.*

### S1B Table. Pairwise association between the factors.

| Pair of factors | Cramér's V / η | Empty cells |
|---|---|---|
| muscle group × tester | 0.14 | 0 of 6 |
| muscle group × participant sex | 0.16 | 1 of 6 |
| muscle group × experiment | 0.46 | 6 of 18 |
| tester × participant sex | 0.56 | 0 of 4 |
| tester × experiment | 0.73 | 3 of 12 |
| participant sex × experiment | 0.74 | 2 of 12 |
| age × muscle group | 0.12 | – |
| age × tester | 0.11 | – |
| age × participant sex | 0.17 | – |
| age × experiment | 0.61 | – |

### S1C Table. Variance inflation in the joint model.

| Factor | df | GVIF | $GVIF^{1/(2\,df)}$ |
|---|---|---|---|
| muscle group | 2 | 2.29 | 1.23 |
| tester | 1 | 2.51 | 1.59 |
| participant sex | 1 | 2.89 | 1.70 |
| age | 1 | 1.65 | 1.28 |
| experiment | 5 | 12.78 | 1.29 |
