## Supplementary material for "From strength to stability: muscular holding capacity characterized by Adaptive Force and its potential relevance for musculoskeletal complaints and non-contact injuries": S2 Table

Schaefer LV, Bittmann FN

### Supplementary Information

**S2 Table. Comparison of the stable and the unstable muscle state within each subgroup.**

Muscle-level values (n = 112 muscles of 68 participants) for every subgroup of the four categorical factors, separately for each AF parameter (S2A–S2E Table), with age covered separately in S2F Table (continuous covariate, no subgroups). Given are the means ± standard deviations of the two muscle states, the mean within-muscle difference MD (unstable − stable; percentage points for AF-Ratio) with its 95% confidence interval, the *p*-value of the paired t-test on MD, and the effect size $d_z$ with its noncentral-t 95% confidence interval. For the three force parameters the relative change, calculated per individual muscle as (unstable − stable)/stable × 100 and averaged across the muscles of the subgroup, is additionally reported; this is the quantity compared between subgroups in Table 2 and Fig 4, for which the MD itself serves in the case of AF-Ratio and $AFosc/AFiso_{max}$. The reference row "All muscles" repeats the overall values of Table 1; its *p*-value is unadjusted.

Two different tests are reported. The *p* of the columns refers to the comparison of the two muscle states within one subgroup and is Holm-adjusted within each parameter across the 13 tests of that table (3 muscle groups, 2 testers, 2 sexes, 6 experiments). All confidence intervals are unadjusted for multiplicity, so an interval excluding zero does not necessarily correspond to a *p*-value below 0.05 after Holm adjustment; this occurs in three subgroups of S2A Table ($AF_{max}$). These within-subgroup tests describe how consistent the difference is across subgroups and are not intended as confirmatory evidence, which is provided by Table 1 for the difference itself and by Table 2 for its modification by the factors.

The *p* of the shaded factor rows refers to the comparison between the subgroups of that factor; it is taken unchanged from Table 2, where Holm is applied within each parameter across the five factors, and is accompanied by the value obtained after exclusion of the pectoralis major. Each adjustment covers the tests actually carried out for the respective question, so the two sets of *p*-values are not interchangeable.

Confidence intervals for $d_z$ are noncentral-t intervals rather than the cluster bootstrap used in Table 1, because subgroups comprise as few as three participants; both methods agree closely where both apply ($AFiso_{max}$, all muscles: noncentral t [−2.20; −1.58] vs. cluster bootstrap [−2.34; −1.61]). Estimates for the pectoralis major rest on three muscles of three participants and are reported for completeness only.

In S2F Table (age), b denotes the slope per year of age from a mixed model with a random intercept for participant, fitted to the same quantity that is compared between subgroups in Table 2 and Fig 4, with *p* Holm-adjusted across the five factors as in Table 2.

$AFiso_{max}$, maximal isometric Adaptive Force; $AF_{max}$, peak Adaptive Force; AFosc, Adaptive Force at the onset of oscillations; AF-Ratio, $AFiso_{max}/AF_{max} \times 100$; b, regression slope; CI, confidence interval; $d_z$, effect size for paired samples; M, mean; MD, mean difference; N, Newton; n, number; *p*, *p*-value; pp, percentage points; SD, standard deviation.

### S2A Table. $AF_{max}$ (N)—peak Adaptive Force.

| | | | Within subgroup: stable vs. unstable | | | | | Compared between subgroups |
|---|---|---|---|---|---|---|---|---|
| **Subgroup** | **n muscles** | **n partici-pants** | **stable M ± SD** | **unstable M ± SD** | **MD [95% CI] (N)** | ***p*** | **$d_z$ [95% CI]** | **relative change [95% CI] (%)** |
| **All muscles** | **112** | **68** | **216.76 ± 65.44** | **230.47 ± 72.84** | **+13.71 [+7.45; +19.96]** | **< 0.001** | **+0.41 [+0.22; +0.60]** | **+7.7 [+3.9; +11.5]** |
| **Muscle group** | | *between subgroups: p = 0.013 without pectoralis major: p = 0.002* | | | | | | |
| elbow flexors | 56 | 48 | 229.45 ± 64.97 | 237.63 ± 73.36 | +8.18 [-0.40; +16.75] | 0.306 | +0.26 [-0.01; +0.52] | +4.4 [-0.6; +9.3] |
| hip flexors | 53 | 51 | 201.54 ± 64.03 | 222.59 ± 73.82 | +21.04 [+11.81; +30.28] | < 0.001 | +0.63 [+0.33; +0.92] | +11.8 [+5.9; +17.8] |
| pectoralis major | 3 | 3 | 248.83 ± 52.99 | 236.25 ± 40.53 | -12.59 [-100.51; +75.34] | 1.000 | -0.36 [-1.50; +0.86] | -3.7 [-37.5; +30.1] |
| **Tester** | | *between subgroups: p = 0.035 without pectoralis major: p = 0.059* | | | | | | |
| tester A | 60 | 36 | 260.84 ± 50.40 | 271.02 ± 67.47 | +10.18 [+2.81; +17.56] | 0.061 | +0.36 [+0.09; +0.62] | +3.2 [+0.3; +6.0] |
| tester B | 52 | 32 | 165.91 ± 38.20 | 183.69 ± 46.00 | +17.78 [+7.17; +28.40] | 0.014 | +0.47 [+0.18; +0.75] | +12.9 [+5.6; +20.2] |
| **Participant sex** | | *between subgroups: p = 1.000 without pectoralis major: p = 0.700* | | | | | | |
| male | 49 | 31 | 276.10 ± 43.20 | 293.77 ± 53.91 | +17.67 [+8.96; +26.37] | 0.002 | +0.58 [+0.28; +0.88] | +6.7 [+3.1; +10.4] |
| female | 63 | 37 | 170.61 ± 35.71 | 181.25 ± 40.25 | +10.63 [+1.68; +19.58] | 0.136 | +0.30 [+0.05; +0.55] | +8.4 [+2.2; +14.6] |
| **Experiment** | | *between subgroups: p = 0.035 without pectoralis major: p = 0.018* | | | | | | |
| Imagery 1 | 10 | 9 | 185.52 ± 64.60 | 196.33 ± 49.37 | +10.81 [-13.61; +35.23] | 1.000 | +0.32 [-0.33; +0.94] | +10.4 [-5.3; +26.2] |
| Imagery 2 | 21 | 12 | 158.39 ± 26.66 | 187.95 ± 31.48 | +29.57 [+12.98; +46.15] | 0.014 | +0.81 [+0.31; +1.30] | +21.7 [+9.6; +33.8] |
| Odorrs | 10 | 9 | 195.75 ± 37.02 | 223.45 ± 31.57 | +27.71 [+5.64; +49.77] | 0.136 | +0.90 [+0.14; +1.62] | +16.1 [+3.3; +28.9] |
| Spindle Procedure | 18 | 11 | 270.75 ± 39.15 | 274.23 ± 51.95 | +3.48 [-9.88; +16.84] | 1.000 | +0.13 [-0.34; +0.59] | +1.1 [-4.0; +6.1] |
| Spindle Manipulation | 23 | 13 | 294.63 ± 33.89 | 322.85 ± 46.62 | +28.22 [+18.40; +38.03] | < 0.001 | +1.24 [+0.69; +1.78] | +9.5 [+6.1; +12.8] |
| Long COVID | 30 | 17 | 182.96 ± 40.68 | 176.89 ± 51.70 | -6.07 [-17.85; +5.70] | 1.000 | -0.19 [-0.55; +0.17] | -3.2 [-10.6; +4.2] |

### S2B Table. AFiso$_{max}$ (N)—maximal isometric AF.

| | | | Within subgroup: stable vs. unstable | | | | | Compared between subgroups |
|---|---|---|---|---|---|---|---|---|
| **Subgroup** | **n muscles** | **n partici-pants** | **stable M ± SD** | **unstable M ± SD** | **MD [95% CI] (N)** | ***p*** | **$d_z$ [95% CI]** | **relative change [95% CI] (%)** |
| **All muscles** | **112** | **68** | **215.36 ± 64.78** | **131.76 ± 62.82** | **-83.59 [-91.86; -75.32]** | **< 0.001** | **-1.89 [-2.20; -1.58]** | **-39.8 [-43.5; -36.1]** |
| **Muscle group** | | *between subgroups: p = 0.019 without pectoralis major: p = 0.083* | | | | | | |
| elbow flexors | 56 | 48 | 227.99 ± 64.06 | 130.84 ± 64.96 | -97.15 [-109.57; -84.74] | < 0.001 | -2.10 [-2.56; -1.62] | -44.0 [-49.1; -38.9] |
| hip flexors | 53 | 51 | 200.11 ± 63.54 | 129.41 ± 61.26 | -70.69 [-80.99; -60.39] | < 0.001 | -1.89 [-2.34; -1.44] | -36.3 [-41.7; -30.9] |
| pectoralis major | 3 | 3 | 248.83 ± 52.99 | 190.46 ± 7.46 | -58.37 [-177.73; +60.98] | 0.170 | -1.21 [-2.74; +0.42] | -21.1 [-63.0; +20.8] |
| **Tester** | | *between subgroups: p = 1.000 without pectoralis major: p = 1.000* | | | | | | |
| tester A | 60 | 36 | 258.81 ± 50.30 | 158.79 ± 63.86 | -100.01 [-111.01; -89.01] | < 0.001 | -2.35 [-2.84; -1.85] | -39.9 [-44.4; -35.3] |
| tester B | 52 | 32 | 165.22 ± 37.72 | 100.57 ± 44.94 | -64.65 [-75.30; -53.99] | < 0.001 | -1.69 [-2.11; -1.26] | -39.6 [-45.8; -33.4] |
| **Participant sex** | | *between subgroups: p = 0.143 without pectoralis major: p = 0.282* | | | | | | |
| male | 49 | 31 | 273.82 ± 43.50 | 180.08 ± 57.16 | -93.74 [-108.15; -79.33] | < 0.001 | -1.87 [-2.33; -1.40] | -34.1 [-39.4; -28.7] |
| female | 63 | 37 | 169.88 ± 35.30 | 94.18 ± 35.43 | -75.70 [-85.12; -66.28] | < 0.001 | -2.02 [-2.45; -1.59] | -44.2 [-49.1; -39.2] |
| **Experiment** | | *between subgroups: p = 0.021 without pectoralis major: p = 0.026* | | | | | | |
| Imagery 1 | 10 | 9 | 184.53 ± 64.70 | 119.49 ± 53.26 | -65.05 [-94.15; -35.95] | 0.001 | -1.60 [-2.54; -0.63] | -34.6 [-48.2; -21.0] |
| Imagery 2 | 21 | 12 | 158.15 ± 26.42 | 102.65 ± 40.93 | -55.50 [-73.69; -37.31] | < 0.001 | -1.39 [-1.98; -0.78] | -35.4 [-46.1; -24.7] |
| Odorrs | 10 | 9 | 193.80 ± 36.03 | 131.43 ± 28.66 | -62.36 [-86.68; -38.04] | < 0.001 | -1.83 [-2.86; -0.78] | -31.1 [-42.0; -20.3] |
| Spindle Procedure | 18 | 11 | 270.27 ± 39.00 | 144.17 ± 66.83 | -126.10 [-155.60; -96.60] | < 0.001 | -2.13 [-2.96; -1.27] | -47.2 [-58.3; -36.1] |
| Spindle Manipulation | 23 | 13 | 290.38 ± 36.75 | 203.45 ± 48.54 | -86.93 [-98.57; -75.28] | < 0.001 | -3.23 [-4.25; -2.19] | -30.5 [-35.2; -25.8] |
| Long COVID | 30 | 17 | 182.39 ± 40.58 | 93.93 ± 45.73 | -88.46 [-99.53; -77.40] | < 0.001 | -2.99 [-3.82; -2.14] | -50.0 [-56.3; -43.6] |

### S2C Table. AF at the onset of oscillations—AFosc (N).

| | | | Within subgroup: stable vs. unstable | | | | | Compared between subgroups |
|---|---|---|---|---|---|---|---|---|
| **Subgroup** | **n muscles** | **n partici-pants** | **stable M ± SD** | **unstable M ± SD** | **MD [95% CI] (N)** | ***p*** | **$d_z$ [95% CI]** | **relative change [95% CI] (%)** |
| **All muscles** | **112** | **68** | **165.71 ± 62.93** | **209.38 ± 66.58** | **+43.67 [+37.46; +49.88]** | **< 0.001** | **+1.32 [+1.06; +1.57]** | **+33.7 [+26.9; +40.6]** |
| **Muscle group** | | *between subgroups: p = 0.035 without pectoralis major: p = 0.015* | | | | | | |
| elbow flexors | 56 | 48 | 186.00 ± 61.43 | 225.45 ± 70.55 | +39.45 [+29.99; +48.92] | < 0.001 | +1.12 [+0.78; +1.45] | +25.6 [+17.3; +33.8] |
| hip flexors | 53 | 51 | 145.17 ± 58.96 | 192.81 ± 60.32 | +47.64 [+39.40; +55.88] | < 0.001 | +1.59 [+1.18; +2.00] | +41.5 [+30.5; +52.5] |
| pectoralis major | 3 | 3 | 149.80 ± 50.36 | 202.10 ± 1.46 | +52.30 [-71.12; +175.71] | 0.210 | +1.05 [-0.49; +2.48] | +48.6 [-103.3; +200.5] |
| **Tester** | | *between subgroups: p = < 0.001 without pectoralis major: p = < 0.001* | | | | | | |
| tester A | 60 | 36 | 208.61 ± 46.99 | 248.21 ± 58.89 | +39.60 [+31.87; +47.33] | < 0.001 | +1.32 [+0.97; +1.67] | +19.5 [+15.4; +23.6] |
| tester B | 52 | 32 | 116.21 ± 37.43 | 164.58 ± 42.61 | +48.37 [+38.26; +58.47] | < 0.001 | +1.33 [+0.95; +1.70] | +50.1 [+37.3; +63.0] |
| **Participant sex** | | *between subgroups: p = 0.337 without pectoralis major: p = 0.291* | | | | | | |
| male | 49 | 31 | 216.56 ± 50.22 | 266.73 ± 48.04 | +50.17 [+40.99; +59.36] | < 0.001 | +1.57 [+1.14; +1.98] | +27.0 [+19.8; +34.3] |
| female | 63 | 37 | 126.16 ± 38.78 | 164.78 ± 39.11 | +38.61 [+30.19; +47.04] | < 0.001 | +1.15 [+0.83; +1.47] | +38.9 [+28.1; +49.8] |
| **Experiment** | | *between subgroups: p = 0.337 without pectoralis major: p = 0.291* | | | | | | |
| Imagery 1 | 10 | 9 | 126.80 ± 47.67 | 175.74 ± 34.15 | +48.94 [+28.65; +69.23] | 0.001 | +1.73 [+0.71; +2.71] | +50.9 [+22.0; +79.8] |
| Imagery 2 | 21 | 12 | 115.81 ± 29.43 | 157.11 ± 33.28 | +41.31 [+23.47; +59.14] | < 0.001 | +1.05 [+0.51; +1.58] | +45.6 [+20.7; +70.4] |
| Odorrs | 10 | 9 | 150.85 ± 37.52 | 206.83 ± 38.17 | +55.97 [+31.22; +80.73] | 0.001 | +1.62 [+0.64; +2.56] | +42.2 [+18.5; +65.8] |
| Spindle Procedure | 18 | 11 | 220.16 ± 42.51 | 263.09 ± 57.34 | +42.93 [+23.72; +62.15] | < 0.001 | +1.11 [+0.51; +1.69] | +20.1 [+11.4; +28.8] |
| Spindle Manipulation | 23 | 13 | 238.16 ± 34.62 | 284.94 ± 33.14 | +46.78 [+39.36; +54.20] | < 0.001 | +2.73 [+1.82; +3.62] | +20.5 [+16.4; +24.5] |
| Long COVID | 30 | 17 | 130.36 ± 41.85 | 167.88 ± 46.47 | +37.53 [+23.96; +51.10] | < 0.001 | +1.03 [+0.58; +1.47] | +35.3 [+20.7; +49.9] |

### S2D Table. AF-Ratio (%)—ratio of $AFiso_{max}$ to $AF_{max}$.

| | | | Within subgroup: stable vs. unstable | | | | |
|---|---|---|---|---|---|---|---|
| **Subgroup** | **n muscles** | **n partici-pants** | **stable M ± SD** | **unstable M ± SD** | **MD [95% CI] (pp)** | ***p*** | **$d_z$ [95% CI]** |
| **All muscles** | **112** | **68** | **99.43 ± 1.81** | **55.78 ± 15.99** | **-43.64 [-46.65; -40.63]** | **< 0.001** | **-2.72 [-3.12; -2.31]** |
| **Muscle group** | | *between subgroups: p = 0.014 without pectoralis major: p = 0.796* | | | | | |
| elbow flexors | 56 | 48 | 99.45 ± 1.39 | 53.39 ± 16.64 | -46.06 [-50.57; -41.54] | < 0.001 | -2.73 [-3.30; -2.16] |
| hip flexors | 53 | 51 | 99.37 ± 2.21 | 56.81 ± 14.20 | -42.55 [-46.46; -38.65] | < 0.001 | -3.01 [-3.64; -2.37] |
| pectoralis major | 3 | 3 | 100.00 ± 0.00 | 82.24 ± 10.46 | -17.76 [-43.74; +8.21] | 0.099 | -1.70 [-3.57; +0.24] |
| **Tester** | | *between subgroups: p = 0.541 without pectoralis major: p = 0.796* | | | | | |
| tester A | 60 | 36 | 99.24 ± 2.31 | 57.62 ± 15.44 | -41.62 [-45.68; -37.56] | < 0.001 | -2.65 [-3.19; -2.11] |
| tester B | 52 | 32 | 99.64 ± 0.90 | 53.67 ± 16.51 | -45.97 [-50.51; -41.43] | < 0.001 | -2.82 [-3.43; -2.21] |
| **Participant sex** | | *between subgroups: p = 0.026 without pectoralis major: p = 0.119* | | | | | |
| male | 49 | 31 | 99.18 ± 2.50 | 61.16 ± 15.99 | -38.02 [-42.66; -33.38] | < 0.001 | -2.35 [-2.90; -1.80] |
| female | 63 | 37 | 99.61 ± 0.95 | 51.60 ± 14.81 | -48.01 [-51.71; -44.31] | < 0.001 | -3.27 [-3.89; -2.64] |
| **Experiment** | | *between subgroups: p = 0.356 without pectoralis major: p = 0.300* | | | | | |
| Imagery 1 | 10 | 9 | 99.53 ± 1.48 | 60.62 ± 19.02 | -38.91 [-51.82; -25.99] | < 0.001 | -2.16 [-3.30; -0.98] |
| Imagery 2 | 21 | 12 | 99.86 ± 0.63 | 54.75 ± 19.66 | -45.12 [-53.99; -36.25] | < 0.001 | -2.32 [-3.14; -1.48] |
| Odorrs | 10 | 9 | 99.08 ± 1.33 | 58.47 ± 8.67 | -40.61 [-46.79; -34.42] | < 0.001 | -4.70 [-6.91; -2.47] |
| Spindle Procedure | 18 | 11 | 99.84 ± 0.52 | 52.04 ± 20.88 | -47.79 [-58.11; -37.47] | < 0.001 | -2.30 [-3.19; -1.40] |
| Spindle Manipulation | 23 | 13 | 98.48 ± 3.51 | 62.63 ± 10.17 | -35.85 [-40.46; -31.23] | < 0.001 | -3.36 [-4.42; -2.29] |
| Long COVID | 30 | 17 | 99.68 ± 0.65 | 51.00 ± 12.73 | -48.68 [-53.48; -43.88] | < 0.001 | -3.79 [-4.82; -2.75] |

### S2E Table. AFosc/AFiso$_{max}$.

| | | | Within subgroup: stable vs. unstable | | | | |
|---|---|---|---|---|---|---|---|
| **Subgroup** | **n muscles** | **n partici-pants** | **stable M ± SD** | **unstable M ± SD** | **MD [95% CI]** | ***p*** | **$d_z$ [95% CI]** |
| **All muscles** | **112** | **68** | **0.756 ± 0.111** | **2.005 ± 1.095** | **+1.250 [+1.042; +1.457]** | **< 0.001** | **+1.13 [+0.89; +1.36]** |
| **Muscle group** | | *between subgroups: p = 0.260 without pectoralis major: p = 0.210* | | | | | |
| elbow flexors | 56 | 48 | 0.806 ± 0.085 | 2.282 ± 1.312 | +1.477 [+1.120; +1.834] | < 0.001 | +1.11 [+0.77; +1.44] |
| hip flexors | 53 | 51 | 0.712 ± 0.110 | 1.765 ± 0.739 | +1.053 [+0.837; +1.270] | < 0.001 | +1.34 [+0.96; +1.71] |
| pectoralis major | 3 | 3 | 0.590 ± 0.085 | 1.063 ± 0.052 | +0.473 [+0.184; +0.763] | 0.020 | +4.06 [+0.38; +7.93] |
| **Tester** | | *between subgroups: p = 1.000 without pectoralis major: p = 1.000* | | | | | |
| tester A | 60 | 36 | 0.805 ± 0.074 | 1.974 ± 1.203 | +1.169 [+0.859; +1.479] | < 0.001 | +0.97 [+0.66; +1.28] |
| tester B | 52 | 32 | 0.698 ± 0.119 | 2.041 ± 0.966 | +1.343 [+1.065; +1.620] | < 0.001 | +1.35 [+0.97; +1.72] |
| **Participant sex** | | *between subgroups: p = 0.555 without pectoralis major: p = 0.804* | | | | | |
| male | 49 | 31 | 0.784 ± 0.095 | 1.854 ± 1.316 | +1.069 [+0.694; +1.445] | < 0.001 | +0.82 [+0.49; +1.14] |
| female | 63 | 37 | 0.733 ± 0.118 | 2.123 ± 0.880 | +1.390 [+1.160; +1.620] | < 0.001 | +1.52 [+1.16; +1.88] |
| **Experiment** | | *between subgroups: p = 0.231 without pectoralis major: p = 0.210* | | | | | |
| Imagery 1 | 10 | 9 | 0.690 ± 0.139 | 1.774 ± 0.700 | +1.084 [+0.604; +1.564] | 0.002 | +1.62 [+0.64; +2.56] |
| Imagery 2 | 21 | 12 | 0.727 ± 0.109 | 1.947 ± 1.186 | +1.221 [+0.669; +1.772] | < 0.001 | +1.01 [+0.47; +1.53] |
| Odorrs | 10 | 9 | 0.774 ± 0.090 | 1.656 ± 0.361 | +0.882 [+0.654; +1.110] | < 0.001 | +2.76 [+1.35; +4.15] |
| Spindle Procedure | 18 | 11 | 0.811 ± 0.075 | 2.613 ± 1.920 | +1.802 [+0.845; +2.758] | 0.002 | +0.94 [+0.37; +1.48] |
| Spindle Manipulation | 23 | 13 | 0.822 ± 0.060 | 1.539 ± 0.478 | +0.717 [+0.509; +0.924] | < 0.001 | +1.50 [+0.89; +2.09] |
| Long COVID | 30 | 17 | 0.707 ± 0.120 | 2.232 ± 0.759 | +1.525 [+1.228; +1.823] | < 0.001 | +1.91 [+1.30; +2.52] |

### S2F Table. Association of age with the change between the muscle states.

| Parameter | b [95% CI] | *p* |
|---|---|---|
| $AF_{max}$ (N) | -0.117 [-0.481; +0.247] % per year | 1.000 |
| $AFiso_{max}$ (N) | -0.031 [-0.372; +0.309] % per year | 1.000 |
| AFosc (N) | +0.497 [-0.115; +1.110] % per year | 0.337 |
| AF-Ratio (%) | +0.046 [-0.223; +0.316] pp per year | 0.738 |
| $AFosc/AFiso_{max}$ | -0.001 [-0.020; +0.017] per year | 1.000 |
