## Supplementary material for "From strength to stability: muscular holding capacity characterized by Adaptive Force and its potential relevance for musculoskeletal complaints and non-contact injuries": S3 Table

Schaefer LV, Bittmann FN

### S3 Table. Sensitivity analyses: the comparison between muscle states on restricted samples.

The main comparison of the difference between muscle states (unstable – stable) for each AF parameter (S3A to S3E) was repeated after omitting each experiment in turn, after omitting subgroups defined by content, for each tester separately, and restricted to the two experiments in which the testers were blinded to the experimental condition. Given are the number of muscles and participants of each restricted sample, the mean within-muscle difference (MD, unstable minus stable), the effect size $d_z$ and the *p*-value of the paired t-test.

*P*-values are unadjusted: these analyses probe the robustness of the pooled result and do not test new hypotheses. The long COVID cohort is the only experiment with diseased participants; its omission is listed under "One experiment omitted". The full sample is given as reference in the first row.

$AFiso_{max}$, maximal isometric Adaptive Force; $AF_{max}$, peak Adaptive Force; AFosc, Adaptive Force at the onset of oscillations; AF-Ratio, $AFiso_{max}/AF_{max} \times 100$; $d_z$, effect size for paired samples; MD, mean difference; N, Newton; n, number; *p*, *p*-value; pp, percentage points; w/o, without.

### S3A. $AF_{max}$ (N)—peak Adaptive Force.

| Analysis | n muscles | n participants | MD (N) | dz | p |
|---|---|---|---|---|---|
| **Full sample** | | | | | |
| all muscles | 112 | 68 | +13.71 | +0.41 | < 0.001 |
| **One experiment omitted** | | | | | |
| w/o Imagery 1 | 102 | 62 | +13.99 | +0.42 | < 0.001 |
| w/o Imagery 2 | 91 | 56 | +10.05 | +0.32 | 0.003 |
| w/o Odors | 102 | 62 | +12.34 | +0.37 | < 0.001 |
| w/o Spindle Procedure | 94 | 57 | +15.67 | +0.46 | < 0.001 |
| w/o Spindle Manipulation | 89 | 55 | +9.96 | +0.29 | 0.008 |
| w/o Long COVID | 82 | 51 | +20.95 | +0.67 | < 0.001 |
| **Subgroups defined by content omitted** | | | | | |
| w/o both spindle experiments (mechanical conditioning) | 71 | 44 | +11.60 | +0.32 | 0.009 |
| w/o imagery and odor experiments (emotionally valent stimuli) | 71 | 41 | +7.46 | +0.24 | 0.048 |
| w/o pectoralis major muscles | 109 | 66 | +14.43 | +0.43 | < 0.001 |
| **Single tester** | | | | | |
| tester A only | 60 | 36 | +10.18 | +0.36 | 0.008 |
| tester B only | 52 | 32 | +17.78 | +0.47 | 0.001 |
| **Testers blinded to the experimental condition** | | | | | |
| Imagery 2 and Odors only | 31 | 21 | +28.97 | +0.85 | < 0.001 |

### S3B. $AFiso_{max}$ (N)—maximal isometric AF.

| Analysis | n muscles | n participants | MD (N) | dz | p |
|---|---|---|---|---|---|
| **Full sample** | | | | | |
| all muscles | 112 | 68 | −83.59 | −1.89 | < 0.001 |
| **One experiment omitted** | | | | | |
| w/o Imagery 1 | 102 | 62 | −85.41 | −1.93 | < 0.001 |
| w/o Imagery 2 | 91 | 56 | −90.08 | −2.11 | < 0.001 |
| w/o Odors | 102 | 62 | −85.67 | −1.92 | < 0.001 |
| w/o Spindle Procedure | 94 | 57 | −75.45 | −2.12 | < 0.001 |
| w/o Spindle Manipulation | 89 | 55 | −82.73 | −1.73 | < 0.001 |
| w/o Long COVID | 82 | 51 | −81.81 | −1.69 | < 0.001 |
| **Subgroups defined by content omitted** | | | | | |
| w/o both spindle experiments (mechanical conditioning) | 71 | 44 | −71.74 | −1.92 | < 0.001 |
| w/o imagery and odor experiments (emotionally valent stimuli) | 71 | 41 | −97.51 | −2.35 | < 0.001 |
| w/o pectoralis major muscles | 109 | 66 | −84.29 | −1.91 | < 0.001 |
| **Single tester** | | | | | |
| tester A only | 60 | 36 | −100.01 | −2.35 | < 0.001 |
| tester B only | 52 | 32 | −64.65 | −1.69 | < 0.001 |
| **Testers blinded to the experimental condition** | | | | | |
| Imagery 2 and Odors only | 31 | 21 | −57.71 | −1.53 | < 0.001 |

### S3C. AFosc (N)—AF at the onset of oscillations.

| Analysis | n muscles | n participants | MD (N) | dz | p |
|---|---|---|---|---|---|
| **Full sample** | | | | | |
| all muscles | 112 | 68 | +43.67 | +1.32 | < 0.001 |
| **One experiment omitted** | | | | | |
| w/o Imagery 1 | 102 | 62 | +43.15 | +1.28 | < 0.001 |
| w/o Imagery 2 | 91 | 56 | +44.22 | +1.39 | < 0.001 |
| w/o Odors | 102 | 62 | +42.46 | +1.29 | < 0.001 |
| w/o Spindle Procedure | 94 | 57 | +43.81 | +1.36 | < 0.001 |
| w/o Spindle Manipulation | 89 | 55 | +42.87 | +1.18 | < 0.001 |
| w/o Long COVID | 82 | 51 | +45.92 | +1.44 | < 0.001 |
| **Subgroups defined by content omitted** | | | | | |
| w/o both spindle experiments (mechanical conditioning) | 71 | 44 | +42.85 | +1.20 | < 0.001 |
| w/o imagery and odor experiments (emotionally valent stimuli) | 71 | 41 | +41.89 | +1.31 | < 0.001 |
| w/o pectoralis major muscles | 109 | 66 | +43.43 | +1.32 | < 0.001 |
| **Single tester** | | | | | |
| tester A only | 60 | 36 | +39.60 | +1.32 | < 0.001 |
| tester B only | 52 | 32 | +48.37 | +1.33 | < 0.001 |
| **Testers blinded to the experimental condition** | | | | | |
| Imagery 2 and Odors only | 31 | 21 | +46.04 | +1.22 | < 0.001 |

### S3D. AF-Ratio (%)—ratio of $AFiso_{max}$ to $AF_{max}$.

| Analysis | n muscles | n participants | MD (pp) | dz | p |
|---|---|---|---|---|---|
| **Full sample** | | | | | |
| all muscles | 112 | 68 | −43.64 | −2.72 | < 0.001 |
| **One experiment omitted** | | | | | |
| w/o Imagery 1 | 102 | 62 | −44.11 | −2.78 | < 0.001 |
| w/o Imagery 2 | 91 | 56 | −43.30 | −2.83 | < 0.001 |
| w/o Odors | 102 | 62 | −43.94 | −2.64 | < 0.001 |
| w/o Spindle Procedure | 94 | 57 | −42.85 | −2.85 | < 0.001 |
| w/o Spindle Manipulation | 89 | 55 | −45.66 | −2.74 | < 0.001 |
| w/o Long COVID | 82 | 51 | −41.80 | −2.49 | < 0.001 |
| **Subgroups defined by content omitted** | | | | | |
| w/o both spindle experiments (mechanical conditioning) | 71 | 44 | −45.11 | −2.90 | < 0.001 |
| w/o imagery and odor experiments (emotionally valent stimuli) | 71 | 41 | −44.30 | −2.84 | < 0.001 |
| w/o pectoralis major muscles | 109 | 66 | −44.35 | −2.84 | < 0.001 |
| **Single tester** | | | | | |
| tester A only | 60 | 36 | −41.62 | −2.65 | < 0.001 |
| tester B only | 52 | 32 | −45.97 | −2.82 | < 0.001 |
| **Testers blinded to the experimental condition** | | | | | |
| Imagery 2 and Odors only | 31 | 21 | −43.66 | −2.61 | < 0.001 |

### S3E. $AFosc/AFiso_{max}$

| Analysis | n muscles | n participants | MD | dz | p |
|---|---|---|---|---|---|
| **Full sample** | | | | | |
| all muscles | 112 | 68 | +1.250 | +1.13 | < 0.001 |
| **One experiment omitted** | | | | | |
| w/o Imagery 1 | 102 | 62 | +1.266 | +1.11 | < 0.001 |
| w/o Imagery 2 | 91 | 56 | +1.256 | +1.15 | < 0.001 |
| w/o Odors | 102 | 62 | +1.286 | +1.12 | < 0.001 |
| w/o Spindle Procedure | 94 | 57 | +1.144 | +1.35 | < 0.001 |
| w/o Spindle Manipulation | 89 | 55 | +1.387 | +1.17 | < 0.001 |
| w/o Long COVID | 82 | 51 | +1.149 | +0.96 | < 0.001 |
| **Subgroups defined by content omitted** | | | | | |
| w/o both spindle experiments (mechanical conditioning) | 71 | 44 | +1.282 | +1.43 | < 0.001 |
| w/o imagery and odor experiments (emotionally valent stimuli) | 71 | 41 | +1.333 | +1.11 | < 0.001 |
| w/o pectoralis major muscles | 109 | 66 | +1.271 | +1.14 | < 0.001 |
| **Single tester** | | | | | |
| tester A only | 60 | 36 | +1.169 | +0.97 | < 0.001 |
| tester B only | 52 | 32 | +1.343 | +1.35 | < 0.001 |
| **Testers blinded to the experimental condition** | | | | | |
| Imagery 2 and Odors only | 31 | 21 | +1.111 | +1.09 | < 0.001 |
