## Supplementary material for "From strength to stability: muscular holding capacity characterized by Adaptive Force and its potential relevance for musculoskeletal complaints and non-contact injuries": S4 Table

$AFiso_{max}$, maximal isometric Adaptive Force; $AF_{max}$, peak Adaptive Force; AFosc, Adaptive Force at the onset of oscillations; AF-Ratio, $AFiso_{max}/AF_{max} \times 100$; df, degrees of freedom of the omnibus contrast; *p*, *p*-value; w/o, without; $\chi^2$, Wald chi-squared statistic.

| Factor | Muscles | $AF_{max}$ (N) | $AFiso_{max}$ (N) | AFosc (N) | AF-Ratio (%) | $AFosc/AFiso_{max}$ |
|---|---|---|---|---|---|---|
| **A. Permutation test** | | | | | | |
| **Muscle group** (df = 2) | all muscles | **0.009 (10.17)** | **0.043 (10.30)** | 0.136 (8.53) | **0.012 (11.36)** | 0.166 (5.49) |
| | w/o pect | **< 0.001 (10.96)** | 0.061 (5.37) | 0.056 (8.02) | 0.738 (1.25) | 0.164 (3.92) |
| **Tester** (df = 1) | all muscles | **0.025 (6.41)** | 1.000 (0.22) | **< 0.001 (15.63)** | 0.539 (1.22) | 1.000 (0.44) |
| | w/o pect | 0.063 (5.28) | 1.000 (0.18) | **< 0.001 (13.52)** | 0.738 (1.04) | 1.000 (0.29) |
| **Participant sex** (df = 1) | all muscles | 1.000 (0.37) | 0.158 (3.78) | 0.323 (2.40) | **0.026 (6.91)** | 0.582 (1.76) |
| | w/o pect | 0.748 (0.18) | 0.287 (2.74) | 0.307 (2.69) | 0.120 (4.87) | 0.879 (1.24) |
| **Age** (df = 1) | all muscles | 1.000 (0.40) | 1.000 (0.03) | 0.323 (2.50) | 0.737 (0.12) | 1.000 (0.02) |
| | w/o pect | 0.748 (0.88) | 1.000 (0.21) | 0.307 (1.20) | 0.799 (0.07) | 1.000 (0.05) |
| **Experiment** (df = 5) | all muscles | 0.057 (13.92) | **0.043 (15.61)** | 0.323 (8.75) | 0.354 (8.70) | 0.147 (11.10) |
| | w/o pect | 0.063 (15.26) | 0.051 (15.67) | 0.307 (9.16) | 0.268 (9.91) | 0.127 (11.31) |

| Factor | Muscles | $AF_{max}$ (N) | $AFiso_{max}$ (N) | AFosc (N) | AF-Ratio (%) | $AFosc/AFiso_{max}$ |
|---|---|---|---|---|---|---|
| **B. Wild cluster bootstrap** | | | | | | |
| Muscle group | all muscles | **0.046** | **0.024** | **0.042** | 0.356 | 0.844 |
| | w/o pect | **0.027** | 0.078 | **0.011** | 0.620 | 0.177 |
| Tester | all muscles | 0.195 | 1.000 | **< 0.001** | 0.496 | 1.000 |
| | w/o pect | 0.229 | 1.000 | **0.002** | 0.620 | 1.000 |
| Participant sex | all muscles | 0.845 | 0.074 | 0.271 | **0.037** | 0.924 |
| | w/o pect | 0.824 | 0.141 | 0.209 | 0.116 | 1.000 |
| Age | all muscles | 0.845 | 1.000 | 0.271 | 0.527 | 1.000 |
| | w/o pect | 0.735 | 1.000 | 0.449 | 0.637 | 1.000 |
| Experiment | all muscles | 0.076 | 0.062 | 0.116 | 0.356 | 0.080 |
| | w/o pect | 0.063 | **0.047** | 0.209 | 0.139 | **0.029** |
